# Spatio-temporal remodeling of extracellular matrix orients epithelial sheet folding

**DOI:** 10.1101/2023.01.13.523870

**Authors:** Alice Tsuboi, Koichi Fujimoto, Takefumi Kondo

## Abstract

Biological systems are inherently noisy; however, they produce highly stereotyped tissue morphology. *Drosophila* pupal wings show a highly stereotypic folding through uniform expansion and subsequent buckling of wing epithelium within a surrounding cuticle sac. The folding pattern produced by buckling is generally stochastic; it is thus unclear how buckling leads to stereotypic tissue folding of the wings. We found that the extracellular matrix (ECM) protein, Dumpy, guides the position and direction of buckling-induced folds. Dumpy anchors the wing epithelium to the overlying cuticle at specific tissue positions. Tissue-wide alterations of Dumpy deposition and degradation yielded different buckling patterns. In summary, we propose that spatio-temporal ECM remodeling shapes stereotyped tissue folding through dynamic interactions between the epithelium and its external structures.

**One-Sentence Summary:** Remodeling of extracellular matrix protein, Dumpy, guides epithelial tissue folding during *Drosophila* wing development.

## Main Text

In origami, a single sheet of paper is folded in a sequence to create a stereotypic shape. Similar folded structures are found everywhere in life, including leaves in buds, insect wings in pupae, and internal organs such as the brain and gastrointestinal tract. Intriguingly, many of the folded structures are highly stereotyped among individuals. These biological folded structures are often formed during development through a physical process called buckling, which generates folds of elastic sheets in response to compressive forces (*1, 2*). Since most of the internal organs develop not in free space but under the physical constraints of the surrounding structures (i.e., neighboring tissues and extracellular matrices [ECM]), tissues need to buckle to relax the compressive stresses imposed by their surroundings. However, unlike origami, folds generated by buckling are generally difficult to predict (*2*) because they are influenced by various physical interactions with their surroundings. Therefore, how tissues achieve stereotypic morphology counteracting the stochasticity of buckling remains an important question.

To address this question, we studied the wing-folding of *Drosophila* pupae. The wing folding process is considered to be caused by buckling with an almost uniform expansion of wing epithelia (fig. S1) in the confined space of the pupal cuticle (brown dotted lines in Fig. 1A) (*3*–*6*). The folding pattern is highly reproducible (fig. S2). We thus focused on the two types of major stereotypic folds, one along four longitudinal veins (L2 –L5) (“folds along veins”, red lines in Fig. 1A) and the other perpendicular to the margin of the wing blade formed by bending the entire wing in the posterior direction (“marginal fold”, blue arrowhead in Fig. 1A). To observe the sequential dynamics of the folding, we performed time-lapse imaging of the pupa (Fig. 1A (i) and B and movie S1). In movie S1, wing expansion and folding along veins initiated from about 39:40 h after puparium formation (APF), which we define as the timing of folding initiation. The wing expansion was not due to cell division, but due to cell surface expansion through cell flattening as reported previously (*4*–*6*) (fig. S3 and Movie S2). Meanwhile, from about 1.7 h after folding initiation (AFI), the marginal fold starts forming at the distal-posterior region (blue arrowhead in Fig. 1A (i)).

**Fig. 1.**
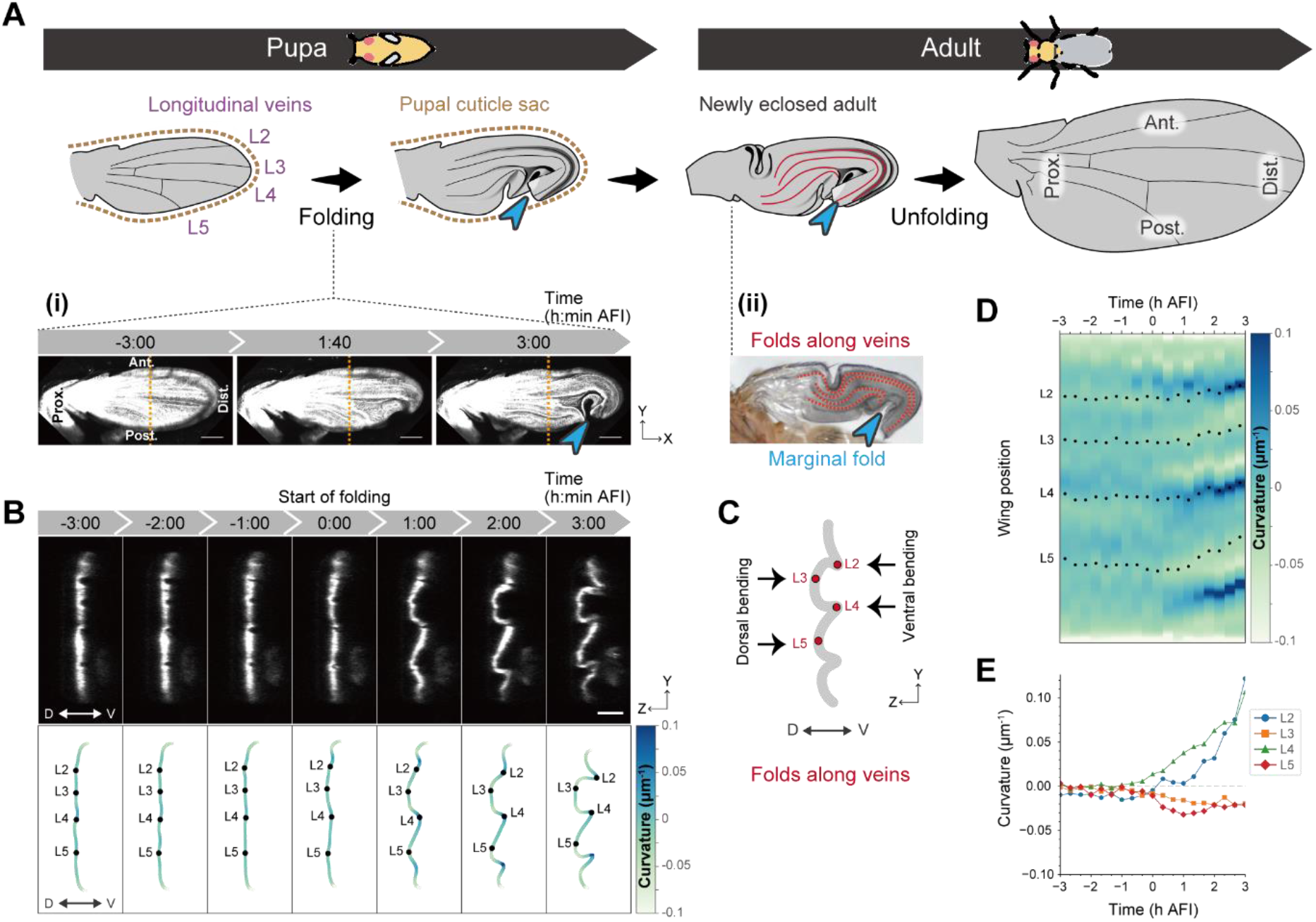
Emergence of stereotypic folding within a cuticle sac. (**A**) Schematics of wing morphogenesis. (i) Maximum projections of confocal time-lapse images in a control wing labelling dorsal cells. (ii) Adult wing just after eclosion has “folds along veins” and “marginal fold.” (**B**) Top: Anterior-posterior cross-sections along the orange dotted lines in (A (i)). Bottom: Color-coded tissue curvature with longitudinal veins (black dots). (**C**) Schematics of folded pupal wing representing “folds along veins” on anterior-posterior cross-section view. (**D, E**) Tissue curvature measurement from a control wing in (B). D: Spatio-temporal color map of curvature with longitudinal veins (black dots). E: Time evolution of the average local curvature around each longitudinal vein. Scale bars, 100 μm (A (i)), 50 μm (B).

We first focused on the “folding along veins” that occurs initially. Among the four folds along veins, two of the longitudinal veins, L2 and L4, bent ventrally (“ Ventral bending”), and the other two, L3 and L5, bent dorsally (“Dorsal bending”) (Fig. 1B upper and C). To map the folding dynamics, we calculated the wing surface curvature along the anterior-posterior axis (Fig. 1B lower). The profiles of the local curvature of five different pupae reproducibly show alternating positive and negative curvatures (Fig. 1B–E and fig. S4A and B). These observations indicate that wings are stereotypically folded along veins.

Since the position and direction of folds produced by buckling are generally stochastic (*2*), there must be unknown mechanisms simultaneously acting to fix the positions and directions of folds in wings. In several developmental processes, such as ventral furrow formation of *Drosophila* gastrula and neural tube formation of vertebrates, epithelial tissues locally generate intrinsic forces using actomyosin for stereotypic folding (*7, 8*). To test whether the actomyosin force is responsible for the stereotypic folding of wings, we knocked down the Myosin II regulatory light chain encoded by *spaghetti squash* (*sqh*) (*9*); however, the wing folding pattern was not affected (fig. S5, Movie S3, and Movie S4), suggesting that the local differences in cell behaviors driven by actomyosin hardly contribute to determining the folding positions in the pupal wing.

We then investigated whether the interactions with the extrinsic environment might control the folding position. We focused on Dumpy (Dpy), a fibrous apical ECM protein that mediates the attachment of the epithelium to the surrounding cuticle in pupa (*10*–*13*). We imaged pre-folded wings of flies harboring the protein-trapped *dpy* allele (*Dpy::YFP*) (*14, 15*) using a multi-photon laser scanning microscope (MP microscope) (Fig. 2A, fig. S6, and Movie S5): We found that Dpy covers the apical cell surface of wing epithelia (Fig. 2 B right “Surface-covering Dpy”) and that Dpy is also present on the pupal cuticle. Additionally, as reported previously (*10, 11*), a dense accumulation of fibrous Dpy in the exuvial space connecting the wing epithelium and the pupal cuticle was found at the wing margin, hinge region, and along the vicinity of L3 and L5 veins (Fig. 2 B left “Exuvial Dpy”). We noticed that the exuvial Dpy along the L3 and L5 veins corresponded to the positions of dorsal bending folds (Fig. 1B–E, fig. S7, and Movie S6). Importantly, the exuvial Dpy along L3 and L5 veins was present both on the dorsal and ventral sides; however, its localization pattern was different. On the dorsal surface, the exuvial Dpy along L3 and L5 veins were continuously localized proximally–distally, whereas on the ventral surface, it was hardly detectable at the distal half of the wing (Fig. 2A (iii), cross-section 3).

**Fig. 2.**
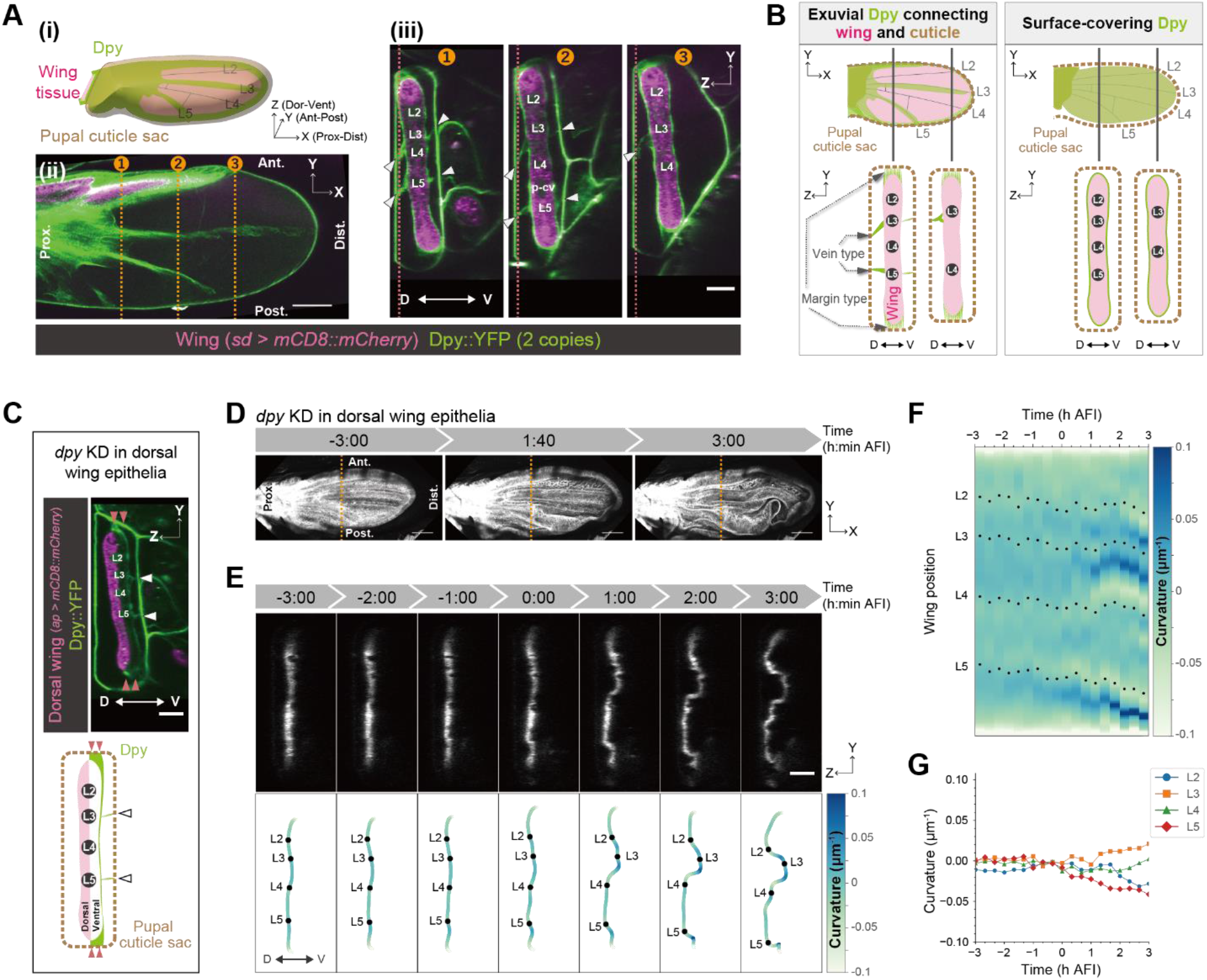
Exuvial Dpy along veins regulates the stereotypic buckling direction. (**A**) (**i**) Schematics of a pre-folded wing. (**ii, iii**) MP microscope snapshot images of a pre-folded control wing which expresses endogenous Dpy::YFP and mCD8::mCherry induced by sd-Gal4 (32.5 h APF). (**ii**) XY view of a slice at the red dotted lines in (iii). (**iii**) Anterior-posterior cross-sections along the orange dotted lines in (ii). White arrowheads indicate exuvial Dpy along veins. (**B**) Schematics showing the two different types of Dpy localization before folding. (**C**) MP microscope snapshot image (top) and schematics (bottom) of a wing with depleted dorsal Dpy. Ventral exuvial Dpy at the margin (red arrowheads) and along veins (white arrowheads) remains. (**D**) Maximum projections of confocal time-lapse images of a wing with depleted dorsal Dpy. (**E**) Top: Anterior-posterior cross-sections along the orange dotted lines in (D). Bottom: Color-coded tissue curvature with longitudinal veins (black dots). (**F, G**) Tissue curvature measurement from the wing in (E). F: Spatio-temporal color map of curvature with longitudinal veins (black dots). G: Time evolution of the average local curvature around each longitudinal vein. Scale bars, 100 μm (A(ii), D), 50 μm (A(iii), C, E).

Based on these observations, we hypothesized that at the positions along L3 and L5 veins, the wing is more strongly anchored to the cuticle dorsally than ventrally, resulting in the dorsally-biased stereotypic bending along L3 and L5 veins during buckling. To test this hypothesis, we depleted Dpy from the dorsal wing (Fig. 2C–G, fig. S8, and Movie S7). In the example shown in Fig. 2E, L2 and L4 veins are bent to the dorsal side, and L3 vein to the ventral side, in the opposite directions to the control. Moreover, the folding position and direction varied among individual wings (Fig. 2F and fig. S4C), and the curvature of most of the veins was close to zero (Fig. 2G and fig. S4D). These results indicate that Dpy depletion from the dorsal surface randomized the folding pattern along longitudinal veins. In contrast, Dpy depletion from the ventral surface exhibited a less significant effect on the folding pattern (fig. S9). Taken together, we conclude that dorsal Dpy, but not ventral Dpy, plays a pivotal role in determining the position and direction of buckling by anchoring the wing epithelium to the cuticle at specific positions in the tissue.

While the wings almost uniformly expand and accumulate compressive stresses in both the anterior-posterior and proximal-distal axes (fig. S1A), folds along veins could relieve the stresses only along the anterior-posterior axis. We thus assume that the compressive stresses not relieved along the proximal-distal axis might drive the forming of the marginal fold. So, we next investigated how the stereotypic marginal fold at the distal-posterior region emerges to relieve the stresses. Then, we closely observed the spatio-temporal dynamics of Dpy distribution in a control wing (Fig. 3A–D and Movie S8) and noticed that Dpy::YFP fluorescent signals on the wing tissue gradually disappeared during folding (see Dpy::YFP signal nearby asterisks in Fig. 3B, B’, and C, and see also purple dotted lines in Fig. 3B’), regardless of its localization type (exuvial vein/margin type, or surface-covering type) (fig. S10, see Fig. 2B for the type of Dpy). Accordingly, the Dpy::YFP fluorescent signal increased at the exuvial space between the cuticle sac and wing tissue (fig. S11), suggesting that Dpy was disassembled and released into the exuvial space during folding. Importantly, the Dpy disappearance starts at the distal-posterior region (see the position of Dpy degradation shown by asterisks in Fig. 3B and C) before forming the marginal fold at the same distal-posterior region (fig. S10D). This implies that the distal-posterior edge of the wing margin started to get detached from the pupal cuticle first (yellow arrowheads in Fig. 3D) and consequently formed the marginal fold (blue arrowhead in Fig. 3D).

**Fig. 3.**
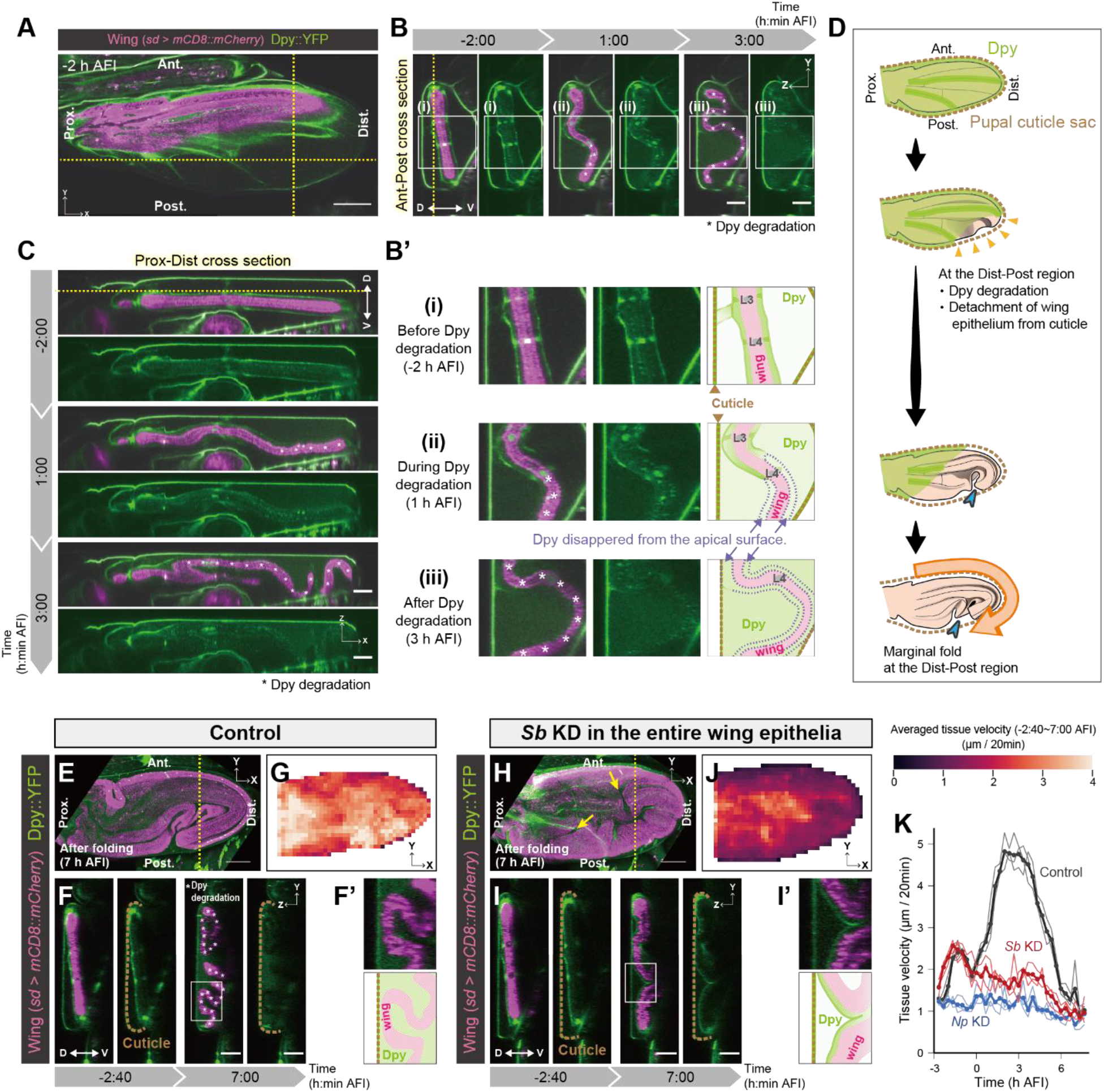
Dpy degradation is indispensable for the marginal fold formation. (**A**–**C**) MP microscope time-lapse images of a control wing expressing mCD8::mCherry induced by sd-Gal4 and endogenous Dpy::YFP. (**A**) XY view of a slice at the yellow dotted lines in (B, C). (**B, C**) Cross-sections along the lines in (A). (**B’**) Magnified views of rectangles in (B) and their schematics. (**D**) Schematics of Dpy degradation and the position of the marginal fold. (**E**–**I**) Confocal time-lapse imaging of Dpy::YFP and PIV analysis in control (E–G) and *Sb* knockdown (H–J). (**E, H**) Maximum projections of post-folded wings (7 h AFI). Yellow arrows in H indicate ectopic folds. (**F, I**) Anterior-posterior cross-sections along the yellow dotted lines in (E) and (H). (**F’, I’**) Magnified views of rectangles in (F, I) and their schematics. (**G, J**) Tissue velocity averaged in each grid throughout the folding process (−2:40–7:00 AFI). (**K**) Temporal dynamics of tissue velocity. N = 3 wings for each genotype (thin lines) and the average (thick lines). Asterisks in B, B’, C and F indicate the wing position where Dpy::YFP signal disappeared from the nearby apical surface. Scale bars, 100 μm (A, E, H), 50 μm (B, C, F, I).

To test whether the starting position of the Dpy degradation determines the position of the marginal fold, we focused on *Stubble* (*Sb*) or *Notopleural* (*Np*), both of which encode transmembrane proteases that are responsible for degrading Dpy (*16*–*19*). When we depleted Sb or Np throughout the wing, the Dpy localization pattern before folding was similar to that in controls (fig. S12); however, we observed abnormalities after folding. The surface-covering and exuvial vein/marginal Dpy proteins remained in their original positions even after the folding was initiated (Fig. 3E–I, fig. S13, fig. S14, Movie S9, and Movie S10), and the Dpy::YFP fluorescent signal at the exuvial space did not increase (Fig. S11), indicating that Sb or Np depletion prevents Dpy disassembly. In terms of the folding process, the Sb- or Np-depleted wings initiated expansion and the first formation of dorsal-bending folds along L3 and L5 veins was in a similar fashion to controls (Movie S9), suggesting that blocking Dpy degradation does not affect the initiation of buckling along veins. However, they failed to eventually form any marginal fold (Fig. 3H and fig. S13A), suggesting that a persistent Dpy anchorage prevents cells from sliding along the wing margin. Further reflecting these abnormalities, their tissue velocities were lower during folding (Fig. 3K, fig. S15A, and Movie S11), especially at the wing margin associated with persistent Dpy anchorage (Fig. 3G and J, fig. S13B, and fig. S15B and C). Moreover, the Sb- or Np-depleted wings formed ectopic folds in the intervein region, some of which crossed over the veins (yellow arrows in Fig. 3H and fig. S13A). Additionally, Sb- or Np-depleted wings showed incomplete cell flattening (fig. S16), suggesting that the persistent Dpy structure also interferes with cell flattening. Taken together, Dpy degradation is indispensable to proceed towards the complete formation of the marginal fold as well as full flattening.

To further examine the impact of Dpy degradation on the marginal fold formation, we sought to alter the Dpy degradation pattern by knocking-down *Sb* in a region-specific manner and examined whether ectopic folds are formed (Fig. 4A–A’’ and fig. S17). Sb depletion in the posterior compartment allows anterior compartment-specific Dpy degradation, which can be regarded as Dpy degradation from the anterior compartment (Fig. 4A’). In this situation, multiple ectopic folds were formed along the anterior margin (Fig. 4A’ and fig. S17E–H). In contrast, Sb depletion in the anterior compartment led to distal-posterior marginal folds similar to those seen in controls (Fig. 4A’’ and fig. S17I and J). Silencing *dpy* expression in such a region-specific manner further supports our hypothesis (Fig. 4B–B’’ and fig. S18): Dpy depletion in the anterior compartment, which could create a situation of premature anterior Dpy degradation before posterior Dpy degradation, resulted in ectopic marginal folds at the distal-anterior position (Fig. 4B’’ and fig. S18G and H). On the other hand, Dpy depletion in the posterior compartment resulted in a distal-posterior marginal fold, located at almost the same position as that of the control (Fig. 4B’ and fig. S18C–F). Taken together, we conclude that spatial coordination of the Dpy degradation specifies the marginal fold position and eventually shapes the higher-order 3D folding.

**Fig. 4.**
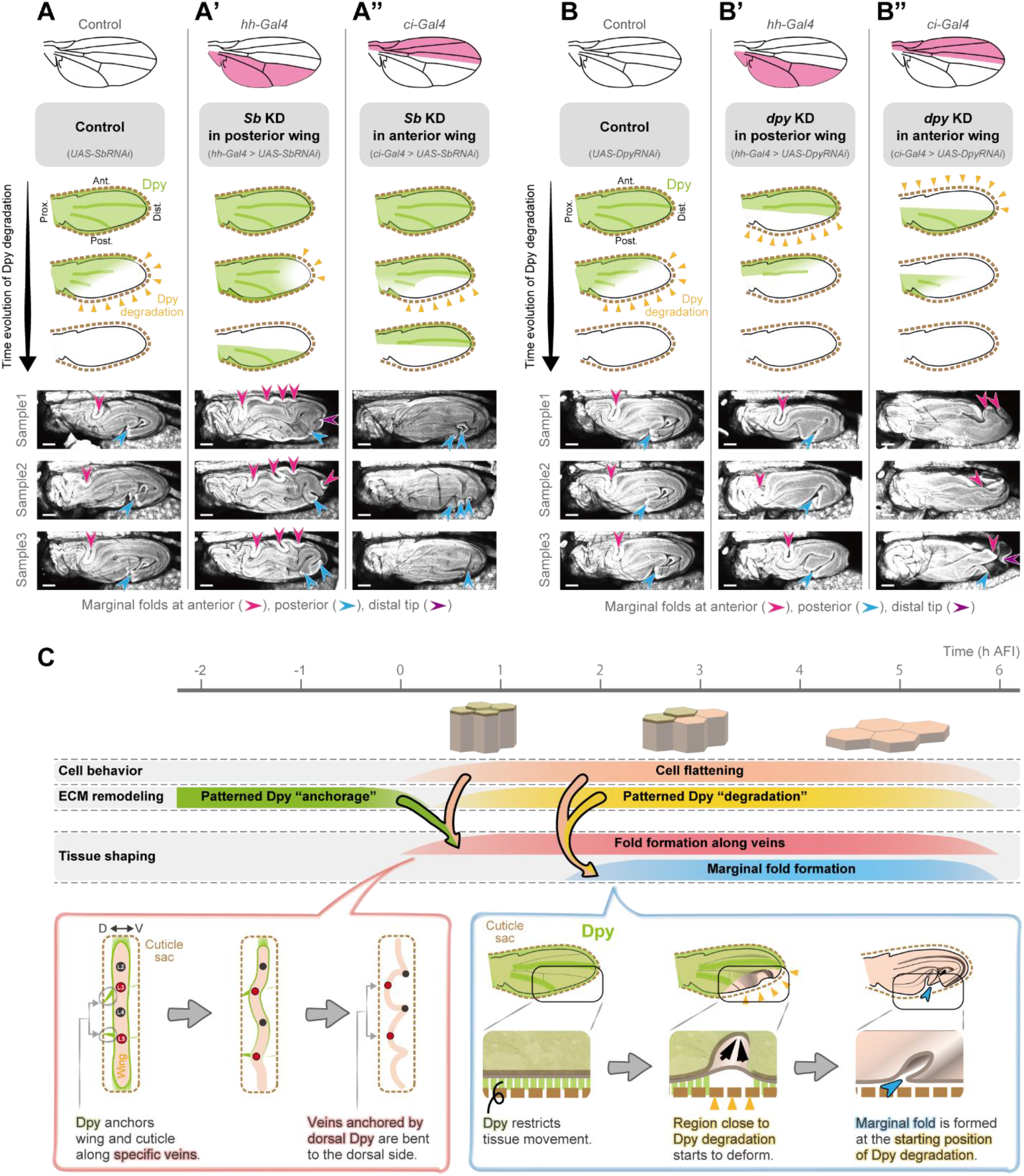
Patterned Dpy degradation specifies the marginal fold position. (**A**–**A’’, B**–**B’’**) Top: Schematics showing the region expressing Gal4 (magenta). Middle: Expected dynamics of Dpy degradation. Yellow arrowheads indicate the starting position of Dpy degradation. Bottom: Maximum projections of confocal snapshot images of GAP43::GFP-expressing-post-folded wings in control (A, B), localized silencing of *Sb* at posterior or anterior region (A’, A’’), localized silencing of *dpy* at posterior or anterior region (B’, B’’). Arrowheads represent folds along the wing margin. Scale bars, 100 μm. (**C**) A model for shaping stereotypic tissue folding.

In summary, we have revealed that spatio-temporally coordinated “deposition” and “destruction” of an ECM protein (Dpy) guides the position and direction of buckling (Fig. 4C). Our results indicate that even though cell populations show spatially homogeneous cellular behaviors (i.e., cell flattening), they can yield stereotypic tissue buckling morphology through the positional information encoded by ECM remodeling. This ECM-based mechanism should confer a potential means to generate diverse and controllable 3D tissue shaping in parallel with cell-intrinsic genetic programming. Future application of this ECM modification to tissue engineering would pave the way for manufacturing precisely folded tissues in any desired manner.

## Acknowledgments

We thank the Kyoto and Bloomington Drosophila Stock Centers, the Vienna Drosophila Resource Center, Magali Suzanne, Robert A. Holmgren, Tetsuya Tabata, Shizue Ohsawa, Yukako Hattori, and Daiki Umetsu for fly stocks; Megumi Hirohata, Kotaro Ikeguchi, Masayo Miki, and Ayumi Nakata for assistance with data analysis and experiments; Housei Wada for the TPIV; Will Draper for the code to calculate curvature; Benoit Aigouy for the Tissue Analyzer and EPySeg; KULIC for imaging with a MP microscope; Shigeo Hayashi, Tadashi Uemura, Tadao Usui, Toshiyuki Harumoto, Miho S. Kitazawa, Satoru Tsugawa, and Daiki Umetsu for comments on the manuscript; and members of the Kondo, Uemura and Fujimoto laboratories for discussions. AT was a JSPS Research Fellow.

## Funding

This work was supported by

JSPS KAKENHI 19J00764 (AT) 21H05779 (AT), 17H06386 (KF), and 16H06280 “ABiS”.

A grant from the NIPPON Genetics (AT)

The Keihanshin Consortium for Fostering the Next Generation of Global Leaders in Research (K-CONNEX) established by the program of Building of Consortia for the Development of Human Resources in Science and Technology, MEXT (TK)

Japan Science and Technology Agency JPMJCR2121 (KF)

## Author contributions

Conceptualization: AT, KF, TK

Formal analysis: AT

Investigation: AT

Funding acquisition: AT, KF, TK

Project administration: AT

Supervision: TK

Writing – original draft: AT, TK

Writing – review & editing: KF

## Data and materials availability

All data are available in the manuscript or the supplementary materials. Raw images and data, image processing macros, codes and excel data for plots are available upon request.

## List of Supplementary Materials

Materials and Methods

Figs. S1 to S18

References (20–31)

Movies S1 to S11

