## Supplementary Materials for "Spatio-temporal remodeling of extracellular matrix orients epithelial sheet folding"

**This PDF file includes:**

Materials and Methods
Figs. S1 to S18
Captions for Movies S1 to S11

**Other Supplementary Materials for this manuscript include the following:**

Movies S1 to S11

**Materials and Methods**

Fly strains

| Reagent type (species) or resource | Designation | Source or reference | Identifiers | Additional information |
|---|---|---|---|---|
| *D. melanogaster* | *w[1118]* | Bloomington Drosophila Stock Center | BDSC: 6326 FlyBase: FBal0018186 | |
| *D. melanogaster* | *ap-GAL4* | Kyoto Stock Center | DGRC: 106798 FlyBase: FBti0002785 | FlyBase symbol: P{GawB}apmd544 |
| *D. melanogaster* | *UAS-Dcr2* | Bloomington Drosophila Stock Center | BDSC: 24651 FlyBase: FBti0100276 | FlyBase symbol: P{UAS-Dcr-2.D}10 |
| *D. melanogaster* | *UAS-DsRed* | Bloomington Drosophila Stock Center | BDSC: 6282 FlyBase: FBti0018001 | FlyBase symbol: P{UAS-AUG-DsRed}A |
| *D. melanogaster* | *sd-Gal4* | Bloomington Drosophila Stock Center | BDSC: 8609 FlyBase: FBti0004225 | FlyBase symbol: P{GawB}sdSG29.1 |
| *D. melanogaster* | *Dpy::YFP* | Kyoto Stock Center | DGRC: 115238 FlyBase: FBti0143891 | FlyBase symbol: PBac{681.P.FSVS-1}dpyCPTI001769 |
| *D. melanogaster* | *UAS-mCD8::mCherry* | Bloomington Drosophila Stock Center | BDSC: 27391 FlyBase: FBti0115768 | FlyBase symbol: P{UAS-mCD8.ChRFP}2 |
| *D. melanogaster* | *UAS-mCD8::mCherry* | Bloomington Drosophila Stock Center | BDSC: 27392 FlyBase: FBti0115769 | FlyBase symbol: P{UAS- |

| | | | | |
|---|---|---|---|---|
| | | | | mCD8.ChRFP}3 |
| *D. melanogaster* | *UAS-dpyRNAi* | Vienna Drosophila Resource Center | VDRC: 44029 FlyBase: FBti0091762 | FlyBase symbol: P{GD4443}v44029 |
| *D. melanogaster* | *UAS-SbRNAi* | Vienna Drosophila Resource Center | VDRC: 1613 FlyBase: FBti0092159 | FlyBase symbol: P{GD473}v1613 |
| *D. melanogaster* | *UAS-NpRNAi* | Vienna Drosophila Resource Center | VDRC: 105297 FlyBase: FBti0121532 | FlyBase symbol: P{KK109020}VIE-260B |
| *D. melanogaster* | *UAS-NpRNAi* | Vienna Drosophila Resource Center | VDRC: 23381 FlyBase: FBti0083222 | FlyBase symbol: P{GD13443}v23381 |
| *D. melanogaster* | *Ubi-GAP43::GFP* | This study | | |
| *D. melanogaster* | *ci-Gal4* | (*20*) | FlyBase: FBal0193963 | FlyBase symbol: P{ci-GAL4.C} |
| *D. melanogaster* | *hh-Gal4* | (*21*) | FlyBase: FBti0017278 | FlyBase symbol: P{GAL4}hhGal4 |
| *D. melanogaster* | *E-Cadherin::GFP* | Bloomington Drosophila Stock Center | BDSC: 60584 FlyBase: FBti0168565 | FlyBase symbol: TI{TI}shg[GFP] |
| *D. melanogaster* | *Act > CD2 > GAL4* | Bloomington Drosophila Stock Center | BDSC: 4780 FlyBase: FBti0012408 | FlyBase symbol: P{GAL4-Act5C(FRT.CD2).P}S |
| *D. melanogaster* | *sqh::eGFP* | (*22*) | FlyBase: | FlyBase |

| | | | | |
|---|---|---|---|---|
| | | | FBti0206965 | symbol: TI{TI}sqh[EGFP.29B] |
| *D. melanogaster* | *Tub-Gal80ts* | Bloomington Drosophila Stock Center | BDSC: 7019 FlyBase: FBti0027796 | FlyBase symbol: P{tubP-GAL80ts}20 |
| *D. melanogaster* | *UAS-Nslmb-vhhGFP4* | Bloomington Drosophila Stock Center | BDSC: 38421 FlyBase: FBti0147362 | FlyBase symbol: P{UAS-Nslmb-vhhGFP4}3 |
| *D. melanogaster* | *ap-lexA* | Bloomington Drosophila Stock Center | BDSC: 54268 FlyBase: FBti0155833 | FlyBase symbol: P{GMR42A06-lexA}attP40 |
| *D. melanogaster* | *LexAop-GAL80* | Bloomington Drosophila Stock Center | BDSC: 32215 FlyBase: FBti0131978 | FlyBase symbol: P{8XLexAop2-IVS-GAL80-WPRE}su(Hw) attP1 |
| *D. melanogaster* | *en-Gal4* | Bloomington Drosophila Stock Center | BDSC: 30564 FlyBase: FBti0003572 | FlyBase symbol: P{en2.4-GAL4}e16E |

Experimental genotypes

**Fig. 1**
**(A (i), B, D, E)** *ap-GAL4/+; UAS-Dcr2/UAS-DsRed*
**(A (ii))** *w[1118]*

**Fig. 2**
**(A)** *sd-Gal4/+; Dpy::YFP, UAS-mCD8::mCherry/Dpy::YFP*
**(C)** *ap-Gal4, Dpy::YFP/UAS-dpyRNAi; UAS-mCD8::mCherry/UAS-Dcr2*
**(D, E, F, G)** *ap-GAL4/UAS-dpyRNAi; UAS-Dcr2/UAS-DsRed*

**Fig. 3**
**(A, B, B', C, E, F, F', G, K Control)** *sd-Gal4/+; Dpy::YFP, UAS-mCD8::mCherry/+*
**(H, I, I', J, K *Sb* KD)** *sd-Gal4/+; Dpy::YFP, UAS-mCD8::mCherry/+; UAS-SbRNAi/+*
**(K *Np* KD)** *sd-Gal4/+; Dpy::YFP, UAS-mCD8::mCherry/UAS-NpRNAi*

**Fig. 4**

**(A)** *Ubi-GAP43::GFP/+; UAS-SbRNAi/+*
**(A')** *Ubi-GAP43::GFP/+; UAS-SbRNAi/hh-Gal4*
**(A'')** *Ubi-GAP43::GFP/+; UAS-SbRNAi/ci-Gal4*
**(B)** *Ubi-GAP43::GFP, UAS-dpyRNAi/+*
**(B')** *Ubi-GAP43::GFP, UAS-dpyRNAi/+; hh-Gal4/+*
**(B'')** *Ubi-GAP43::GFP, UAS-dpyRNAi/+; ci-Gal4/+*

**Fig. S1**
**(A Control, B Control, C Control)** *ap-GAL4/+; UAS-Dcr2/UAS-DsRed*
**(A *dpy* KD, B *dpy* KD , C *dpy* KD)** *ap-GAL4/UAS-dpyRNAi; UAS-Dcr2/UAS-DsRed*

**Fig. S2**
*w[1118]*

**Fig. S3**
**(B, C)** *y, w, hs-flp; E-Cadherin::GFP; Act > CD2 > GAL4, UAS-mCD8::mCherry*

**Fig. S4**
**(A, B)** *ap-GAL4/+; UAS-Dcr2/UAS-DsRed*
**(C, D)** *ap-GAL4/UAS-dpyRNAi; UAS-Dcr2/UAS-DsRed*

**Fig. S5**
**(A, C bottom)** *y, w, sqh::eGFP/Y; Tub-Gal80ts/+; UAS-Nslmb-vhhGFP4, UAS-mCD8::mCherry/ci-Gal4*
**(B, D bottom)** *y, w, sqh::eGFP/Y; Tub-Gal80ts/+; UAS-Nslmb-vhhGFP4, UAS-mCD8::mCherry/hh-Gal4*
**(C top)** *y, w, sqh::eGFP/+; Tub-Gal80ts/+; UAS-Nslmb-vhhGFP4, UAS-mCD8::mCherry/ci-Gal4*
**(C middle)** *y, w, sqh::eGFP/Y; Tub-Gal80ts/+; ci-Gal4/TM3, Sb ftz-lacZ*
**(D top)** *y, w, sqh::eGFP/+; Tub-Gal80ts/+; UAS-Nslmb-vhhGFP4, UAS-mCD8::mCherry/hh-Gal4*
**(D middle)** *y, w, sqh::eGFP/Y; Tub-Gal80ts/+; hh-Gal4/TM3, Sb ftz-lacZ*

**Fig. S6**
**(A, B, C)** *sd-Gal4/+; Dpy::YFP, UAS-mCD8::mCherry/Dpy::YFP*

**Fig. S7**
*sd-Gal4/+; Dpy::YFP, UAS-mCD8::mCherry/+*

**Fig. S8**
**(A)** *ap-Gal4, Dpy::YFP/+; UAS-mCD8::mCherry/UAS-Dcr2*
**(B)** *ap-Gal4, Dpy::YFP/UAS-dpyRNAi; UAS-mCD8::mCherry/UAS-Dcr2*

**Fig. S9**
**(A)** *sd-Gal4/+; ap-lexA, UAS-mCD8::mCherry/Dpy::YFP; UAS-Dcr2, LexAop-GAL80/+*
**(B, C, D, E)** *sd-Gal4/+; ap-lexA, UAS-mCD8::mCherry/+; UAS-Dcr2, LexAop-GAL80/+*

**(F)** *sd-Gal4/+; ap-lexA, UAS-mCD8::mCherry/Dpy::YFP, UAS-dpyRNAi; UAS-Dcr2, LexAop-GAL80/+*
**(G, H, I, J)** *sd-Gal4/+; ap-lexA, UAS-mCD8::mCherry/UAS-dpyRNAi; UAS-Dcr2, LexAop-GAL80/+*

**Fig. S10**
**(A, B, C, C', D, D')** *sd-Gal4/+; Dpy::YFP, UAS-mCD8::mCherry/+*

**Fig. S11**
**(A Control, B Control)** *sd-Gal4/+; Dpy::YFP, UAS-mCD8::mCherry/+*
**(A *Sb* KD, B *Sb* KD )** *sd-Gal4/+; Dpy::YFP, UAS-mCD8::mCherry/+; UAS-SbRNAi/+*
**(A *Np* KD, B *Np* KD )** *sd-Gal4/+; Dpy::YFP, UAS-mCD8::mCherry/UAS-NpRNAi*

**Fig. S12**
**(A)** *sd-Gal4/+; Dpy::YFP, UAS-mCD8::mCherry/+; UAS-SbRNAi/+*
**(B)** *sd-Gal4/+; Dpy::YFP, UAS-mCD8::mCherry/UAS-NpRNAi*

**Fig. S13**
**(A, B, C, C')** *sd-Gal4/+; Dpy::YFP, UAS-mCD8::mCherry/UAS-NpRNAi*

**Fig. S14**
**(A)** *sd-Gal4/+; Dpy::YFP, UAS-mCD8::mCherry/+*
**(B)** *sd-Gal4/+; Dpy::YFP, UAS-mCD8::mCherry/+; UAS-SbRNAi/+*
**(C)** *sd-Gal4/+; Dpy::YFP, UAS-mCD8::mCherry/UAS-NpRNAi*

**Fig. S15**
**(A Control, B Control, C Control)** *sd-Gal4/+; Dpy::YFP, UAS-mCD8::mCherry/+*
**(A *Sb* KD, B *Sb* KD , C *Sb* KD )** *sd-Gal4/+; Dpy::YFP, UAS-mCD8::mCherry/+; UAS-SbRNAi/+*
**(A *Np* KD, B *Np* KD , C *Np* KD )** *sd-Gal4/+; Dpy::YFP, UAS-mCD8::mCherry/UAS-NpRNAi*

**Fig. S16**
**(A Control, B Control)** *sd-Gal4/+; UAS-mCD8::mCherry/E-Cadherin::GFP*
**(A *Sb* KD, B *Sb* KD )** *sd-Gal4/+; UAS-mCD8::mCherry/E-Cadherin::GFP; UAS-SbRNAi/+*
**(A *Np* KD, B *Np* KD )** *sd-Gal4/+; UAS-mCD8::mCherry/E-Cadherin::GFP; UAS-NpRNAi/+*

**Fig. S17**
**(B)** *Ubi-GAP43::GFP/+; UAS-SbRNAi/+*
**(C)** *sd-Gal4/+; Ubi-GAP43::GFP/+; +/UAS-DsRed*
**(D)** *sd-Gal4/+; Ubi-GAP43::GFP/+; UAS-SbRNAi/+*
**(E)** *Ubi-GAP43::GFP/+; hh-Gal4/UAS-DsRed*
**(F)** *Ubi-GAP43::GFP/+; UAS-SbRNAi/hh-Gal4*
**(G)** *Ubi-GAP43::GFP/en-Gal4; +/UAS-DsRed*
**(H)** *Ubi-GAP43::GFP/en-Gal4; UAS-SbRNAi/+*
**(I)** *Ubi-GAP43::GFP/+; ci-Gal4/UAS-DsRed*
**(J)** *Ubi-GAP43::GFP/+; UAS-SbRNAi/ci-Gal4*

**Fig. S18**
**(B)** *Ubi-GAP43::GFP, UAS-dpyRNAi/+*
**(C)** *Ubi-GAP43::GFP/+; hh-Gal4/UAS-DsRed*
**(D)** *Ubi-GAP43::GFP, UAS-dpyRNAi/+; hh-Gal4/+*
**(E)** *Ubi-GAP43::GFP/en-Gal4; +/UAS-DsRed*
**(F)** *Ubi-GAP43::GFP, UAS-dpyRNAi/en-Gal4*
**(G)** *Ubi-GAP43::GFP/+; ci-Gal4/UAS-DsRed*
**(H)** *Ubi-GAP43::GFP, UAS-dpyRNAi/+; ci-Gal4/+*

**Movie S1.**
*ap-GAL4/+; UAS-Dcr2/UAS-DsRed*

**Movie S2.**
*y, w, hs-flp; E-Cadherin::GFP; Act > CD2 > GAL4, UAS-mCD8::mCherry*

**Movie S3.**
*y, w, sqh::eGFP/Y; Tub-Gal80ts/+; UAS-Nslmb-vhhGFP4,* UAS-mCD8::mCherry*/ci-Gal4*

**Movie S4.**
*y, w, sqh::eGFP/Y; Tub-Gal80ts/+; UAS-Nslmb-vhhGFP4,* UAS-mCD8::mCherry*/hh-Gal4*

**Movie S5.**
*sd-Gal4/+; Dpy::YFP, UAS-mCD8::mCherry/Dpy::YFP*

**Movie S6.**
*sd-Gal4/+; Dpy::YFP, UAS-mCD8::mCherry/+*

**Movie S7.**
*ap-GAL4/UAS-dpyRNAi; UAS-Dcr2/UAS-DsRed*

**Movie S8.**
*sd-Gal4/+; Dpy::YFP, UAS-mCD8::mCherry/+*

**Movie S9.**
**(Control)** *sd-Gal4/+; Dpy::YFP, UAS-mCD8::mCherry/+*
**(*Sb* KD)** *sd-Gal4/+; Dpy::YFP, UAS-mCD8::mCherry/+; UAS-SbRNAi/+*
**(*Np* KD)** *sd-Gal4/+; Dpy::YFP, UAS-mCD8::mCherry/UAS-NpRNAi*

**Movie S10.**
**(Control)** *sd-Gal4/+; Dpy::YFP, UAS-mCD8::mCherry/+*
**(*Sb* KD)** *sd-Gal4/+; Dpy::YFP, UAS-mCD8::mCherry/+; UAS-SbRNAi/+*
**(*Np* KD)** *sd-Gal4/+; Dpy::YFP, UAS-mCD8::mCherry/UAS-NpRNAi*

**Movie S11.**
**(Control)** *sd-Gal4/+; Dpy::YFP, UAS-mCD8::mCherry/+*

**(*Sb* KD)** *sd-Gal4/+; Dpy::YFP, UAS-mCD8::mCherry/+; UAS-SbRNAi/+*
**(*Np* KD)** *sd-Gal4/+; Dpy::YFP, UAS-mCD8::mCherry/UAS-NpRNAi*

Husbandry conditions
Fly stocks were kept in vials at 18 or 25°C with a standard corn-sugar-yeast food and transferred every 4 or 3 weeks, respectively. For cross experiments, flies were flipped every 3–4 days and vials containing offspring were raised at 18, 25, or 29°C until dissection or imaging.

Live imaging
Pupae for live imaging were prepared as previously described with some modifications (*23*, *24*): White prepupae (WPP) were collected and kept at 25°C (unless otherwise noted) until they reached the appropriate stage. Staged pupae were washed with water, dried on KimWipe, and then mounted on a glass slide using double-sided tape with the right wing on top. The pupal case was carefully removed with forceps so as not to cause an influx of hemolymph into the wings. Filter paper (ADVANTEC 02103020) wetted with water was placed around the pupa to avoid desiccation. A bank of silicone oil compounds (Shin-Etsu Chemical Co., Ltd., HIVAC-G) was placed around the dissected pupa and covered with a coverslip. The coverslip was carefully placed parallel to the dorsal wing cuticle to avoid shear forces, which cause abnormal wing packing. We note that dissection before folding for time-lapse imaging reproducibly formed the stereotypic folds along veins and margin at the distal-posterior region, however, sometimes caused ectopic folds at the anterior wing margin. Therefore, we dissected wings after the folding was completed and confirmed that the positions of marginal fold in control wings are reproducible among individuals (fig. S17B (n=10/10 samples) and fig. S18B (n=10/10 samples)). For Fig. 2C, Fig. 3E to K, fig. S5, fig. S8, fig. S10, fig. S13, fig. S14, fig. S15, and fig. S16, exposed wings were covered with a small drop of halocarbon oil (a 4:1 mixture of halocarbon oil 700 and 27, Sigma Aldrich). Images were obtained at room temperature (at 22–25°C) using Zeiss LSM 710 upright laser confocal microscope with a 20× objective (EC Plan-Neofluar 20x/0.50 M27), LSM 800 inverted laser confocal microscope with a 10× objective (Plan-Apochromat 10×/0.45 M27), a 20× objective (Plan-Apochromat 20×/0.8 M27), and a silicon oil immersion objective (LD LCI Plan-Apochromat 40×/1.2 Imm Korr DIC M27), or Olympus FV1000MPE-IX83 MP microscope (Spectra-Physics InSight DeepSee Laser set to 930 nm wavelength) with a silicon oil immersion objective (UPLSAPO30XS). For the time-lapse recording, each wing was imaged every 20 min except for fig. S16 (every 1 h). For the protease knockdown experiment in the entire wings (Fig. 3H–K, fig. S11, fig. S12, fig. S13, fig. S14B and C, fig. S15, fig. S16, and fig. S17D), we excluded pupae whose pre-folded wings are abnormal contour shape (small and round) from the live imaging analysis because the abnormal shape makes it difficult to evaluate the impact of the proteases on wing folding.

Image processing
All images were processed using Fiji software (https://fiji.sc/). The range of intensity was adjusted with the "setMinAndMax(min, max)" function by setting the minimum and maximum displayed pixel values. For Fig. 1A (i, ii), B, D, and E, Fig. 2A and C–G, Fig. 3A–C and E–K, fig S1A Adult, B, and C, fig. S2, fig. S4, fig. S5, fig. S6, fig. S8, fig. S9, fig. S10, fig. S11, fig. S12, fig. S13, fig. S14, and fig. S15, images were rotated once to bring the anterior side on top and the distal side to the right. In case the imaging was started before the wings were folded, the image was rotated so that the line connecting the intersection of L2 and L3 veins and the distal end point of L3 vein is

horizontal to the XY axis of the image. Stack projection views were generated by applying maximum intensity Z-projection. Cross-sections of a stack were generated by using the "*Image/Stacks/Reslice*" command. For Fig. 2A and C, Fig. 3A–C, fig. S1A Adult, B, and C, fig. S6, fig. S8, and fig. S9A and F, tiles spanning a wing were stitched together using Pairwise Stitching or Grid/Collection Stitching Plugin. For images captured by a MP microscope (Olympus FV1000MPE-IX83), the stitched images were smoothed by using the "Process/Smooth" command. The positions of veins are inferred from the absence of a fluorescent signal within the tissue. For 3D view images rotating around the y-axis in Movie S10, Fiji 3D Viewer plugin was used.

Characterization and curvature measurement of folds along four longitudinal veins

Cross-sections of a stack of time-lapse images were generated along the anterior-posterior axis that passes through the intersection of the L4 longitudinal vein and the posterior cross vein at 38 h after puparium formation (APF). Wing surface outlines were extracted based on DsRed or mCD8::mCherry signals using Fiji and analyzed using custom-written code in Python. In Fiji, the wing surface contour was manually traced from the cross-section images (YZT images) using the segmented line tool, interpolated with a width of 0.5 pixels using the "*Edit/Selection/Interpolate*" command. After that, we acquired a series of wing surface contour points (x, y). The position of longitudinal veins (L2, L3, L4, L5) was determined from the original stacked images (XYZT images) and projected to the corresponding position in the cross-section images because it was difficult to recognize the vein positions in cross-section images of wings labeling only dorsal or ventral surface. In Python, we fit a third-order polynomial to a sliding window along the wing surface contour points $(x, y)$ and measured the first and second derivatives of position with respect to distance along the contour, from which we calculated local signed curvature $k$ using the following formula:

$$k = (x'y'' - y'x'')/(x'^2 + y'^2)^{3/2}.$$

In our analysis, the curve bending to the ventral side is defined as positive. We evaluated the curvature at a series of 200 points equally spaced in distance along the contour. Of the 200 points that constitute the wing surface, the points closest to the vein coordinates were defined as the locations of the vein. The curvature values were color-coded, and the vein positions were represented by black dots on the corresponding coordinates in sequential time series images (Fig. 1B, Fig. 2E, fig. S4A and C, and fig. S9C, D, H and I) or on the spatio-temporal heat maps (Fig. 1D, Fig. 2F, fig. S4A and C, and fig. S9D and I). Curvature values of 11 points in the vicinity of the vein were averaged and plotted as a function of time (Fig. 1E, Fig. 2G, fig. S4B and D, and fig. S9E and J). Python code for calculating curvature was downloaded from the Harvard Dataverse [Draper, W. E. Analysis code. Harvard Dataverse. doi:10.7910/DVN/CHOTF0 (2016).49.] which was published by Draper and Liphardt (*25*).

Particle image velocimetry (PIV) analysis

Time-lapse images were rotated, maximum intensity Z-projections were applied (see Image processing for details), and background signals from outside wing tissue were removed from the images by erasing the area outside the Region of Interests (ROIs) that were manually located along the wing margin ("*Edit/Clear Outside*" command in Fiji). PIV analysis was performed on the RFP channels (sd-Gal4>UAS-mCD8::mCherry) of the processed time-lapse images using TPIV customized version of ver 3.3 (https://signaling.riken.jp/tools/imagej-plugins/490/, a Fiji plugin developed by Housei Wada). Parameters for TPIV were set as follows: Window width and height (pixel): 50, Overlap (%): 50, Extended value: 3, SubPixel value: 10, Color Method: Absolute, Max length limit: 16, Check “OverWrite,” “WithOriginalImage,” and “CorrectByMedian”. In brief, images were split into 50×50-pixel (31.2×31.2 μm) interrogation windows, and the interrogation window overlap was set to 50% to calculate tissue velocities (respective displacement vectors between subsequent frames). Each tissue velocity was color-coded according to its magnitude by colorizing each vector into 8 colors using “Absolute” mode with 16 pixels (9.98 μm) as the maximum value (Fig. S15A). The tissue velocity output data from TPIV was further analyzed with custom-written code in Python and plotted using Python or Excel. For the temporal dynamics of tissue velocity (Fig. 3K), the magnitude of all the tissue velocities within the wing margin was averaged for each time frame and plotted as a function of time. For the spatial color maps of tissue velocity (Fig. 3, G and J, fig. S13B and fig. S15B), the temporal average of all velocity vector magnitudes throughout the folding process (-2:40–7:00 AFI) was calculated within wing margin in each grid [25×25 pixel (15.6×15.6 μm)] which was compartmentalized according to the location of the initial point of each vector. For the spatio-temporal tissue velocity (Fig. S15C), we first spatially categorized the tissue velocity vectors within the wing margin according to the normalized distance to wing margin = floor($D_{v\text{-}m}$ / G), with $D_{v\text{-}m}$ being the minimum distance from the initial point of each vector to the wing margin and G being the constant value representing the distance between adjacent vectors [25×25 pixel (15.6×15.6 μm)]. The spatio-temporal tissue velocity was calculated by averaging the tissue velocities for each spatial and temporal category, and a heatmap was plotted as a function of time (h AFI) (*x*-axis) and normalized distance to wing margin (*y*-axis). Only if the number of PIV vectors belonging to the category was ten or more, were they displayed in the heat map.

### Plasmid Construction and transgenesis for *Ubi-GAP43::GFP* strain

GFP fused with an N-terminal 20-amino-acid peptide of plasma membrane localization tag from growth-associated protein 43 (GAP43) gene was amplified by PCR with PrimeSTAR (Takara Bio) and primers (5′-AACAGATCTGCGGCCGCAACATGCTGTGCTGTATGCGAAGAACCAAAC AGGTTGAAAAAAATGATGAGGACCAAAAGATTATGAGTAAAGGAGAAGAACTTTTC -3′ (Underline corresponds to plasma membrane localization tag) and 5´-GGATTCCTCCACGGGGTACCTTATTTGTATAGTTCATCCATGCCATGTG-3´) and subcloned into pUbi-attB (*26*) with the In-Fusion PCR cloning kit (Clontech). With this plasmid, a transgenic strain was generated by the phiC31 integrase-mediated transgene integration into an attP target site of the ZH-22A (*27*).

### Observation of adult wings using a stereo microscope

Adult flies right after eclosion were immediately transferred to the freezer until they stopped moving. Images were taken using an Olympus SZ61 stereo microscope equipped with a LUMIX DMC-G7H digital camera.

### Quantification of wing tissue area

To prevent overcrowding, 15 females and 3 males were crossed in a vial at 25°C and transferred to a new vial with fresh food every 3–4 days. For the preparation of pupal wings, pupae before wing folding (28–34 h APF) were dissected as described in the “Live imaging” section above. For the preparation of adult wings, adult flies that had eclosed at least 24 h ago, which is sufficient time for wing expansion in the wild type, were transferred into 1.5 ml tubes or dishes (IWAKI non-treated 35-mm culture dishes 1000-035, IWAKI, Shizuoka, Japan) and moved to the freezer. Adult wings were cut off, washed in 70% EtOH, and mounted on a glass slide in 80% glycerol in 1×PBS. Both pupal and adult wings were observed at room temperature (25°C) using LSM 800 inverted laser confocal microscope with Plan-Apochromat 10×/0.45 M27. We applied maximum intensity Z-projection. In the case of adult wings, tiles of the maximum intensity Z-projections spanning a wing were stitched together using the Pairwise Stitching Plugin in Fiji. The wing area was measured by manually tracing the wing outline from subcostal break to alula notch.

Comparison of planer contour shape of adult and pupal wing

For comparison of the planer contour shape of pupal and adult wings in the right panel of fig. S1A, the pupal wing image was colored orange, and the adult wing image was colored blue. Then, the images were made translucent, sized so that the pupal and adult wings were approximately the same size, rotated, and overlayed in Adobe Illustrator.

Quantification of cell area

For fig. S3B and C, larvae were incubated in a water bath at 34°C for 30 min to induce clones of mCherry-marked cells. Then, 1, 25, or 26 h after heat shock, white prepupae (WPP) were collected and kept at 25°C until the appropriate stage. For fig. S16, control (*sd-Gal4>UAS-mCD8::mCherry*), *Sb* knockdown (*sd-Gal4>UAS-mCD8::mCherry, UAS-SbRNAi*), and *Np* knockdown (*sd-Gal4>UAS-mCD8::mCherry, UAS-NpRNAi*) wings expressing E-Cadherin::GFP were raised at 25°C. Time-lapse imaging was performed using LSM800 (equipped with LD LCI Plan-Apochromat 40×/1.2 Imm Korr DIC M27) as described above. Since it is difficult to measure the cell area in the deep folded regions, images cropped from the region where the tissue is relatively flat were used for cell area quantification. Based on the adherens junction marker E-Cadherin::GFP, we segmented images and tracked cells at the intervein region using Fiji plugin Tissue Analyzer (*28*, *29*). The brief segmentation procedure is as follows:

- Apply maximum intensity Z-projection to 7 slices (1 μm per slice) containing an adherens junction marker E-Cadherin::GFP, apply median filter (1-pixel radius), convert to RGB color, and save as image sequence (TIF format) with Fiji.
- Automated segmentation images were obtained using Tissue Analyzer or the python package EPySeg (*30*). In the case of Tissue Analyzer, select the channel of E-Cadherin::GFP and click the "Detect bonds V3 (save watershed)" in the "Segmentation" tab. In the case of EPySeg, select a pre-trained model (2D epithelial segmentation) in the "Model" tab and set the "Channel of interest (COI)" to the channel of E-Cadherin::GFP and output segmentation images in "Tissue Analyzer mode" in the "Predict" tab. (All of the following procedures were conducted using Tissue analyzer.)
- Manually correct the mistakes of the automated segmentation and remove cells in and around the vein region ("Correction" tab).
- Click "Finish all" and "Check finish all" in the "PostProcess" tab. The parameter (4-way vertex vs bond cut-off) was set to 2.
- Click "Autocenter based on 2D correction" in the "Recenter" tab.
- Click "Track cells (dynamics tissue)" in the "Tracking" tab. For fig. S3C, mCherry-marked cells were used as landmarks to check for tracking errors (e.g., cell swapping errors, cell pairing errors), and if any errors were found, they were manually fixed from the edit button "Correct/edit cell tracks" to track the same cells through time. Click "Check track" and "Update track mask" to finalize cell tracking.

- Click “Generate/Update database” in the “SQLite DB” tab. The parameter (4-way vertex vs bond cut-off) was set to 2.
- Click “Export cell data” in the “Plots” tab to save the area of cells in .csv format.

For fig. S3C, the same cells were tracked over time using the location of the clonally mCherry-marked cells as landmarks, and the cell area for each cell was plotted as a function of time. For fig. S16B, the area of cells at the same time (h AFI) was averaged and plotted as a function of time.

<u>NSlmb-vhhGFP4 mediated degradation of Sqh::eGFP</u>
To deplete Sqh from the stage right before wing folding, deGradFP (*9*) and Gal80ts (*31*) systems were used in combination. In Sqh::eGFP knock-in flies harboring Tub-Gal80ts, NSlmb-vhhGFP4 and mCD8::mCherry were overexpressed by ci-Gal4/hh-Gal4 driver with temperature shift from 18°C to 29°C at 48 h APF. Here, 48 h APF at 18°C approximately corresponds to about 21–22 h APF at 25°C (*32*). For fig. S5A and B, temperature-shifted pupae were kept at 29°C for about 4 h in the incubator to select pupae with the necessary markers for observation using a fluorescent stereomicroscope, and time-lapse imaging of the degradation process of Sqh::eGFP was performed with 20 min interval as described above, except that the temperature was changed using a thermal controller (TOKAI HIT, Thermo Plate, TPi-SQH26FT). Note that the thermal controller was set to 31°C to keep samples between 29°C and 30°C at room temperature (25°C). For fig. S5C and D, temperature-shifted pupae were kept at 29°C for 1 day in an incubator and then dissected to take snapshot images of folded wings. Female pupae harboring both Sqh::GFP and Sqh (endogenous non-GFP-fused Sqh which is not subject to degradation) and male pupae without driving NSlmb-vhhGFP4 mediated knockdown were used as control, and male pupae driving NSlmb-vhhGFP4 mediated knockdown were used as Sqh knockdown. Female and male pupae were distinguished at the WPP stage by identifying the enlarged male gonads. Observed pupae were raised while mounted on the cover glass, and their sexes were reconfirmed at the adult stage.

<u>Quantification of intensity</u>
For fig. S10C’ and D’ and fig. S14, images for quantifying the signal intensities were maximum projections (fig. S10C’ and D’ and fig. S14A (i), B (ii), and C (iii)) or cross-sections of a stack (fig. S14A (iv), B (v), and C (vi)) processed with Fiji as described in the section “Image processing”. Intensity profiles along the straight line (line width: 10 pixels) were obtained using the “*Analyze/Plot Profile*” command in Fiji. For fig. S10C’ and D’, the intensity profiles were normalized by the maximum intensity value among -1:40–1:40 AFI (40-min increments).

For fig. S11, we obtained the images using the same microscope setup and generated cross-sections of a stack processed with Fiji as described in the section “Image processing.” In the cross-section image, ROIs [10×10 pixels (6.2×6.2 μm)] were set at exuvial space between the dorsal cuticle and dorsal wing surface to measure the averaged Dpy::YFP intensity in the ROIs. mCD8::mCherry signal in wing tissue was used to make sure the position of the wing surface.

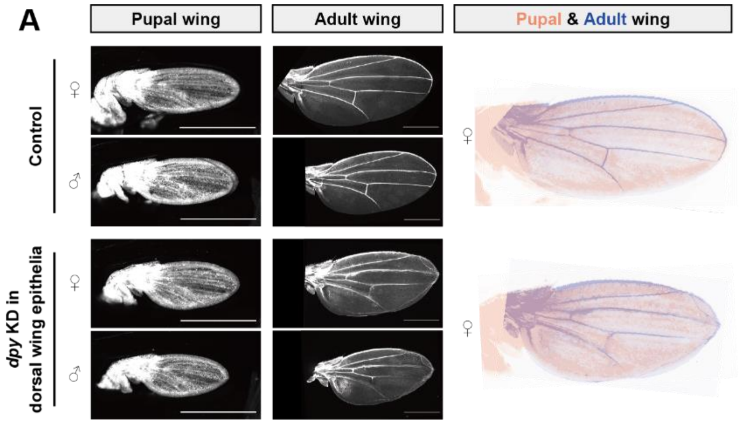


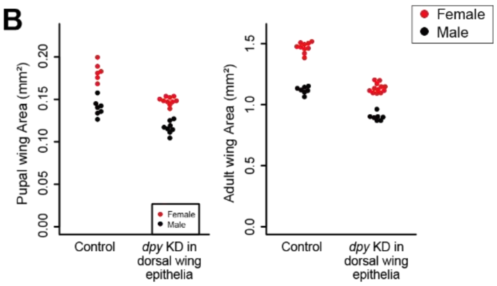


C

| | Wing area ratio (Adult/Pupa) |
|---|---|
| Control ♀ | 8.0 |
| Control ♂ | 8.0 |
| *dpy* KD in dorsal wing epithelia ♀ | 7.7 |
| *dpy* KD in dorsal wing epithelia ♂ | 7.7 |

**Fig. S1. Wing tissue almost uniformly expands from the pupal stage before folding to the adult stage.**

**(A)** Pupal and adult wings in control (*ap-Gal4>UAS-DsRed, UAS-Dcr2*) and *dpy* knockdown at the dorsal surface (*ap-Gal4>UAS-dpyRNAi, UAS-DsRed, UAS-Dcr2*). Pupal wings before folding (left) appear to be miniature adult wings (middle), as evidenced by the almost exact match of the planer contours when a pupal wing and an adult wing are overlapped (right). Pupal and adult wings in the right panel are the same as the female wings in the left and middle panels. While the wings with depleted dorsal Dpy were slightly smaller than the control (see also (B)), probably due to the halving of the marginal Dpy, the overall contour of the wing appears to be kept almost normal before folding.

**(B)** Wing area of pupal and adult wings in control (*ap-Gal4>UAS-DsRed, UAS-Dcr2*) and *dpy* knockdown at the dorsal surface (*ap-Gal4>UAS-dpyRNAi, UAS-DsRed, UAS-Dcr2*). N = 6 wings (female) and N = 7 wings (male) in control at the pupal stage, N = 10 wings (female) and N = 8 wings (male) in *dpy* knockdown at the pupal stage, N = 10 wings (female) and N = 7 wings (male) in control at the adult stage, and N = 12 wings (female) and N = 7 wings (male) in *dpy* knockdown at the adult stage.

**(C)** Wing area ratio (Adult/Pupa) calculated from data shown in B.

Scale bars, 500 µm (A).

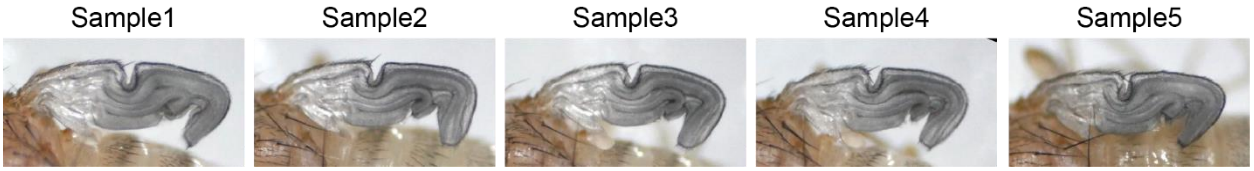


**Fig. S2. Wing folding structure right after eclosion is reproducible among individuals.** Stereotypic wing folding of control (*w[1118]*) adult flies just after eclosion. Each figure indicates a separate individual. The original image of Sample1 is the same as the inset of Fig. 1A (ii).

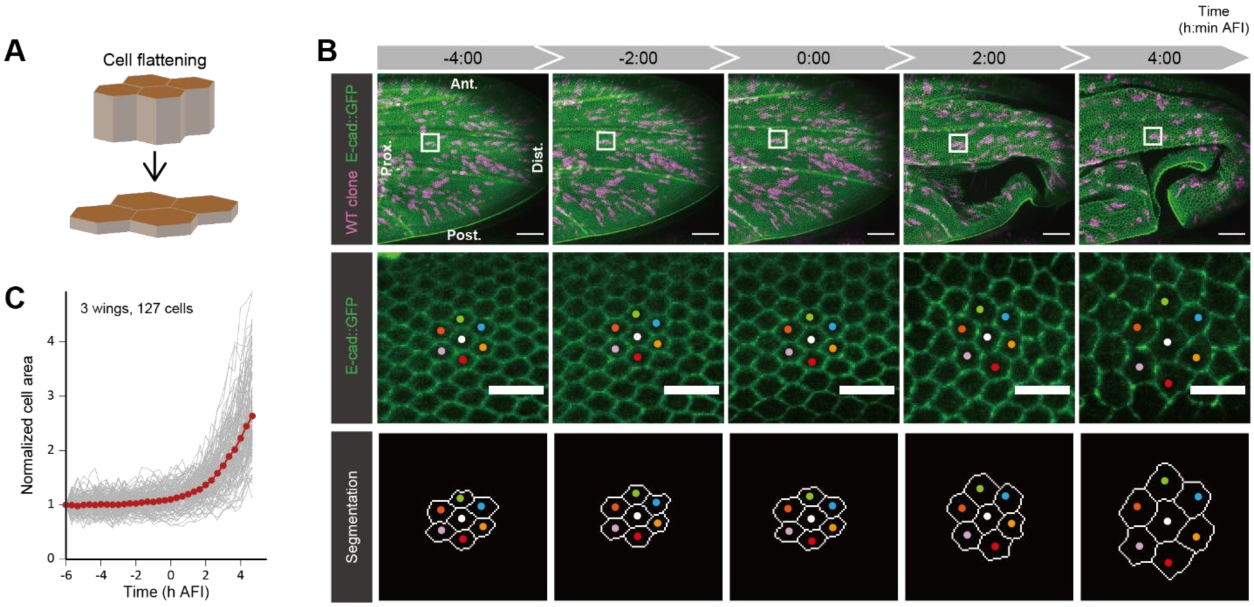


**Fig. S3. Apical area of wing epithelial cells increases during wing folding.**

**(A)** Schematics represent cell flattening during wing folding. Apical side of cells is colored brown, and their lateral side is colored light-brown. Note that the schematics are simplified representations of the cell shape, and that the actual cells have hair and pedestals (the bulge at the base of hairs) on the apical surface.

**(B)** Top: Maximum projections of confocal time-lapse images in the control wing expressing E-Cadherin::GFP with clonally mCherry-marked cells, which can be used as landmarks to track the same cells over time. Middle and bottom: Magnified views of squares in the top panels (middle) and their segmented images based on junction marker E-Cadherin::GFP (bottom). Dots labeled in a specific color represent tracked cells.

**(C)** Apical cell area normalized at -6 h AFI as a function of time (h AFI). Gray lines are individual cell data (N = 127 cells from 3 wings), and the red line is the average. The area of the apical cell surface barely changes until about 0 h AFI, but thereafter it increases rapidly and reaches 2.6-fold by 4:40 AFI.

Scale bars, 50 µm (B top) and 10 µm (B middle).

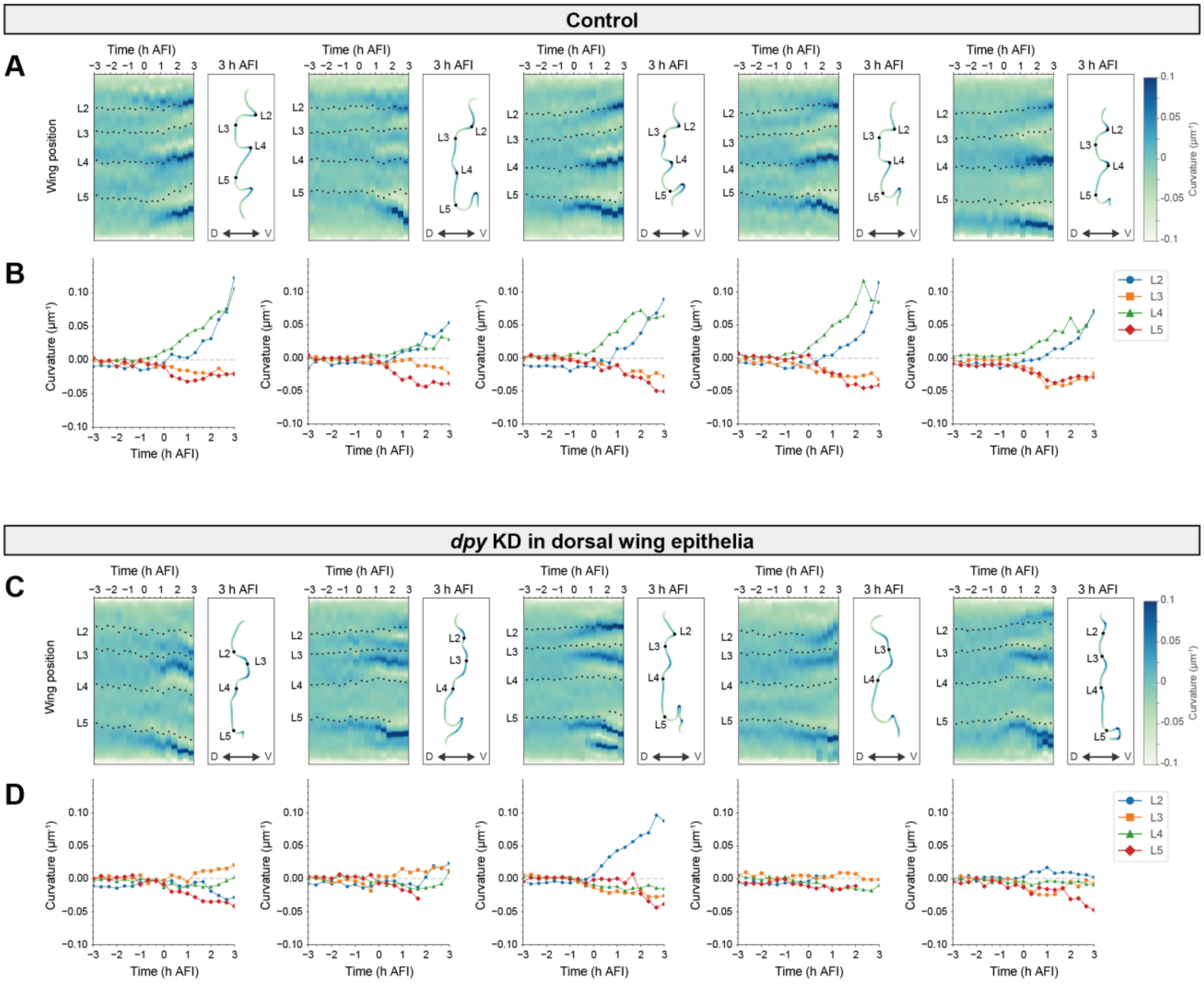


**Fig. S4. Quantification of wing folding structure in control wings and wings with depleted dorsal Dpy.**

**(A–D)** Tissue curvature measurement from time-lapse imaging of five control wings (*ap-Gal4>UAS-DsRed*, *UAS-Dcr2*) (A, B) and five wings with depleted dorsal Dpy (*ap-Gal4>UAS-dpyRNAi*, *UAS-DsRed*, *UAS-Dcr2*) (C, D). Left-most figures are the same as Fig. 1D and E for control, and Fig. 2F and G for *dpy* knockdown. A, C: Spatio-temporal color maps of curvature (left). Anterior-posterior cross-sections showing tissue curvature at 3 h AFI (right). Black dots represent the positions of L2, L3, L4, and L5 longitudinal veins. Note that some of the positions of longitudinal veins after folding were not shown because the veins cannot be recognized due to the folded structure. B, D: Time evolution of local curvature averaged around longitudinal veins. Five different control wings show similar concavity and convexity patterns, in which L2 and L4 longitudinal veins show positive curvature and L3 and L5 veins show negative curvature. In contrast, five wings with depleted dorsal Dpy show varied concavity and convexity patterns, and most of the curvature at the veins was close to zero.

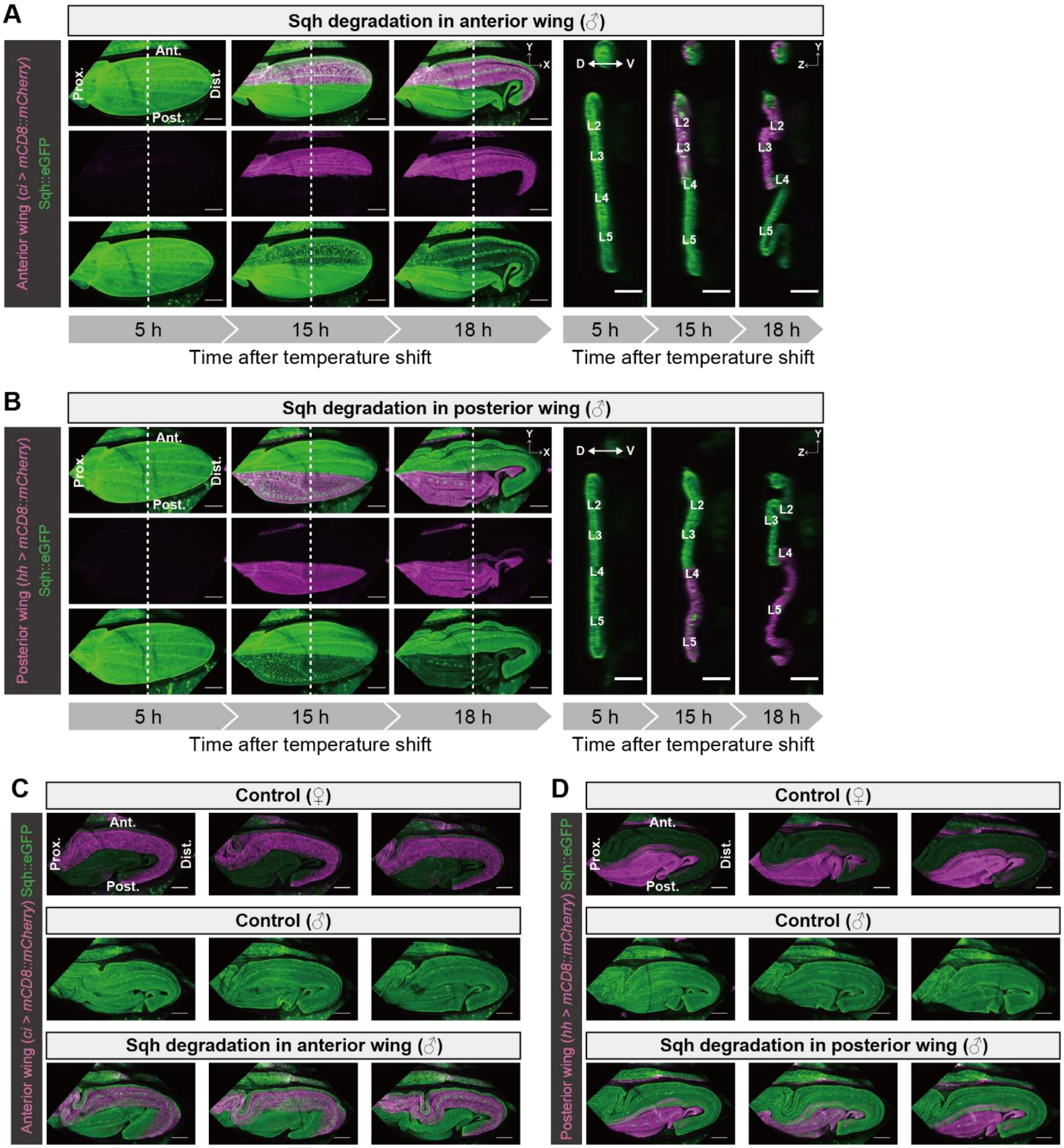


**Fig. S5. Nanobody-based knockdown of Myosin II regulatory light chain.**

**(A–D)** NSlmb-vhhGFP4 mediated knockdown of Sqh::eGFP at the anterior region (A, C) or posterior region (B, D). A, B: Confocal time-lapse images of the Sqh::eGFP degradation process in which time (h) after the temperature shift is shown at the bottom. XY views are the maximum projections and YZ views are the anterior-posterior cross-sections along the white dotted lines in XY views. C, D: Maximum projections of confocal snapshot images of wings after folding in control and Sqh::eGFP knockdown. Female pupae harboring both Sqh::GFP and Sqh (endogenous non-GFP-fused Sqh) (*y, w, sqh::eGFP/+; Tub-Gal80ts/+; UAS-Nslmb-vhhGFP4, UAS-mCD8::mCherry/ci-Gal4* (C top), *y, w, sqh::eGFP/+; Tub-Gal80ts/+; UAS-Nslmb-vhhGFP4,*

*UAS-mCD8::mCherry/hh-Gal4* (D top)) and male pupae without driving NSlmb-vhhGFP4 mediated knockdown (*y, w, sqh::eGFP/Y; Tub-Gal80ts/+; ci-Gal4/TM3, Sb ftz-lacZ* (C middle), *y, w, sqh::eGFP/Y; Tub-Gal80ts/+; hh-Gal4/TM3, Sb ftz-lacZ* (D middle)) were considered as control, and male pupae driving NSlmb-vhhGFP4 mediated knockdown (*y, w, sqh::eGFP/Y; Tub-Gal80ts/+; UAS-Nslmb-vhhGFP4, UAS-mCD8::mCherry/ci-Gal4* (A, C bottom), *y, w, sqh::eGFP/Y; Tub-Gal80ts/+; UAS-Nslmb-vhhGFP4, UAS-mCD8::mCherry/hh-Gal4* (B, D bottom)) were considered as Sqh knockdown.
Scale bars, 100 μm (XY views of A and B, C, D) and 50 μm (YZ views of A and B).

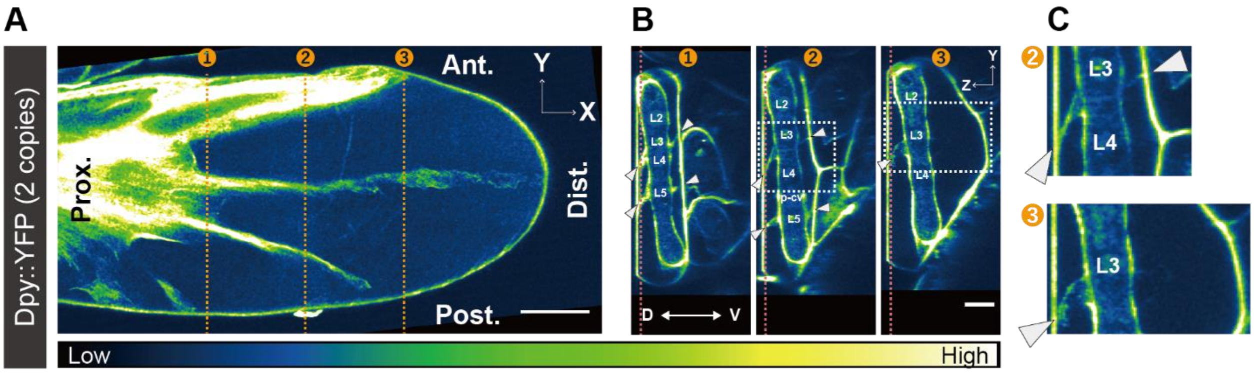


**Fig. S6. Dpy localization on the wing before folding.**

**(A, B)** Dpy::YFP signal in Fig. 2A (ii, iii) was color-coded depending on its intensity. A: XY view of a slice between the dorsal cuticle and wing surface at the position of the red dotted lines in (B). B: Anterior-posterior cross-sections along the orange dotted lines in (A).

**(C)** Magnified view of a rectangle in (B). White arrowheads indicate exuvial Dpy along veins between the wing and cuticle. Note that exuvial Dpy is present both in panel 2 and panel 3 on the dorsal side but not in panel 3 on the ventral side.

Scale bars, 100 µm (A), 50 µm (B).

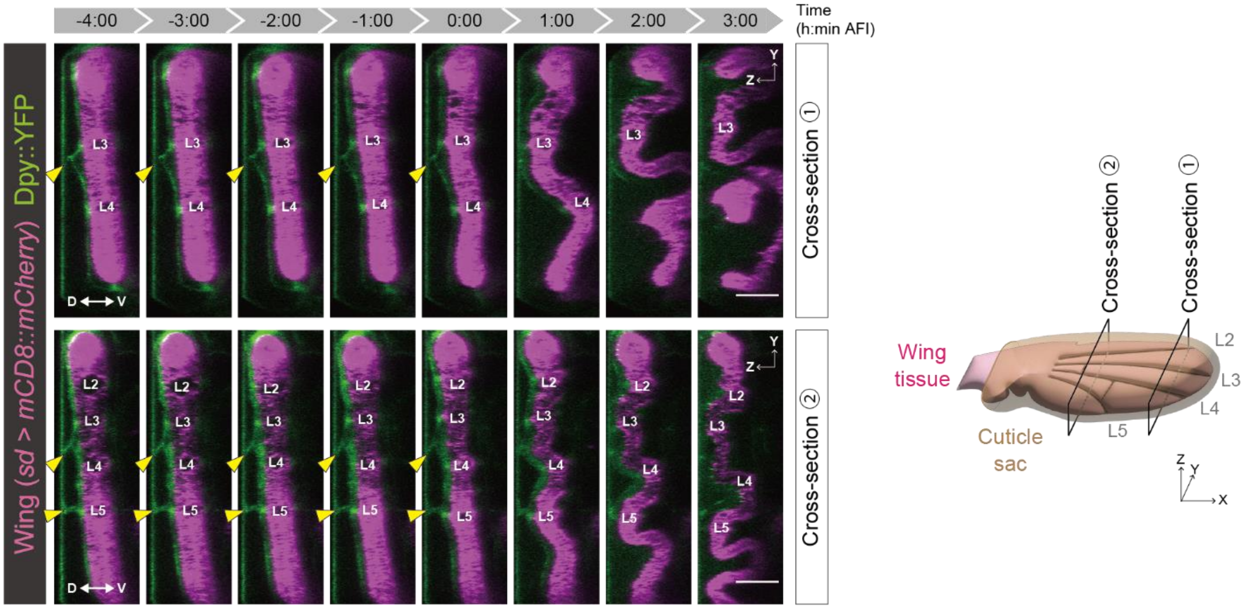


**Fig. S7. Exuvial Dpy along the L3 and L5 veins corresponds to the positions of dorsal bending folds.**

Anterior-posterior cross-sections of confocal time-lapse images in a control wing expressing mCD8::mCherry induced by sd-Gal4 and endogenous Dpy::YFP. The wing positions where exuvial Dpy is localized (yellow arrowheads) were bent to the dorsal side.
Scale bars, 50 µm.

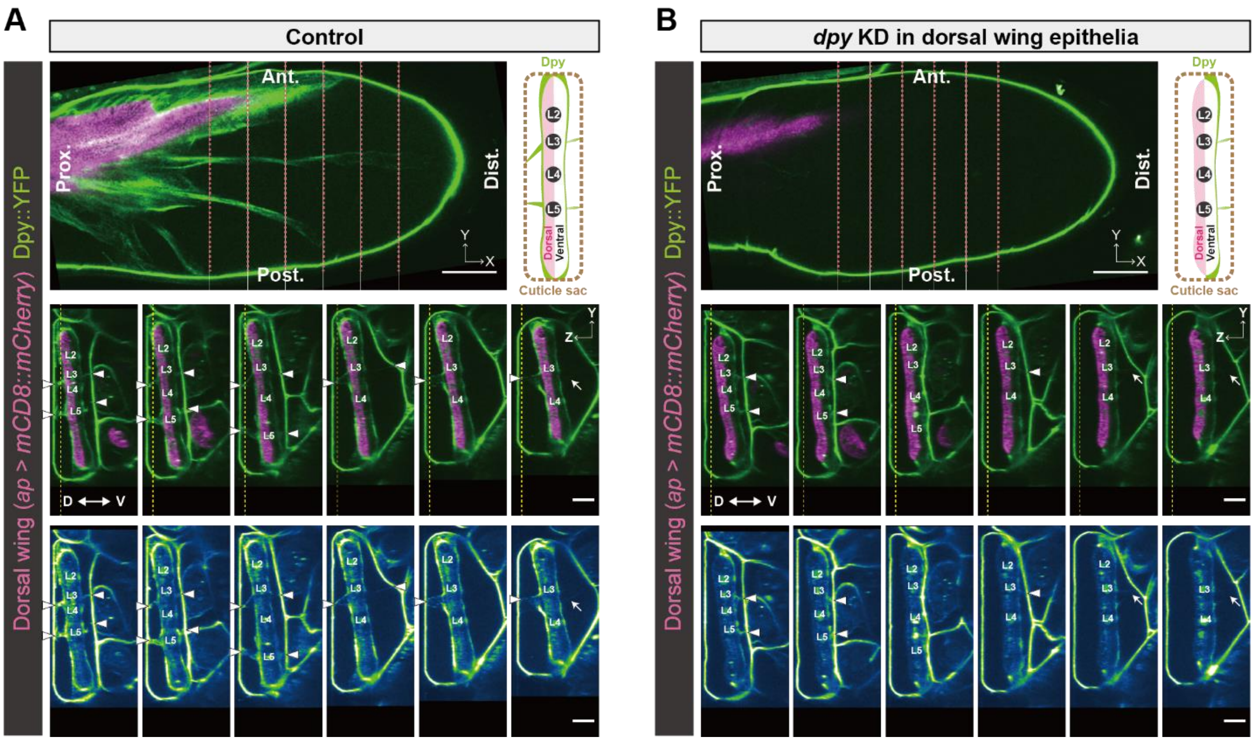


**Fig. S8. Dpy localization in control wings and wings with depleted dorsal Dpy.**

**(A, B)** MP microscope snapshot images of wings expressing Dpy::YFP before folding (33:00 APF (A), 33:05 APF (B)) in a control wing (*ap-Gal4>UAS- mCD8::mCherry, UAS-Dcr2*) (A) and a wing depleted with dorsal Dpy (*ap-Gal4>UAS-dpyRNAi, UAS- mCD8::mCherry, UAS-Dcr2*) (B). Both wings express mCD8::mCherry to label dorsal cells. Top left: XY views of a slice between the dorsal cuticle and wing surface. Top right: Schematics of representative Dpy (green) distribution on wings labeling dorsal cells (magenta) inside a cuticle sac (brown). Bottom: Anterior-posterior cross-sections along the red lines in the top panels. White arrowheads indicate exuvial Dpy along veins between the wing and cuticle, and white arrows indicate a very weak exuvial Dpy signal on the ventral side. Yellow dotted lines represent the corresponding Z slice position of the top panel. Left-most cross-section of (B) is the same as Fig. 2C. In a wing with depleted dorsal Dpy, Dpy was effectively diminished on the dorsal side, but the ventral side preserved its Dpy expression.

Scale bars, 100 μm (XY views of A and B), 50 μm (YZ views of A and B).

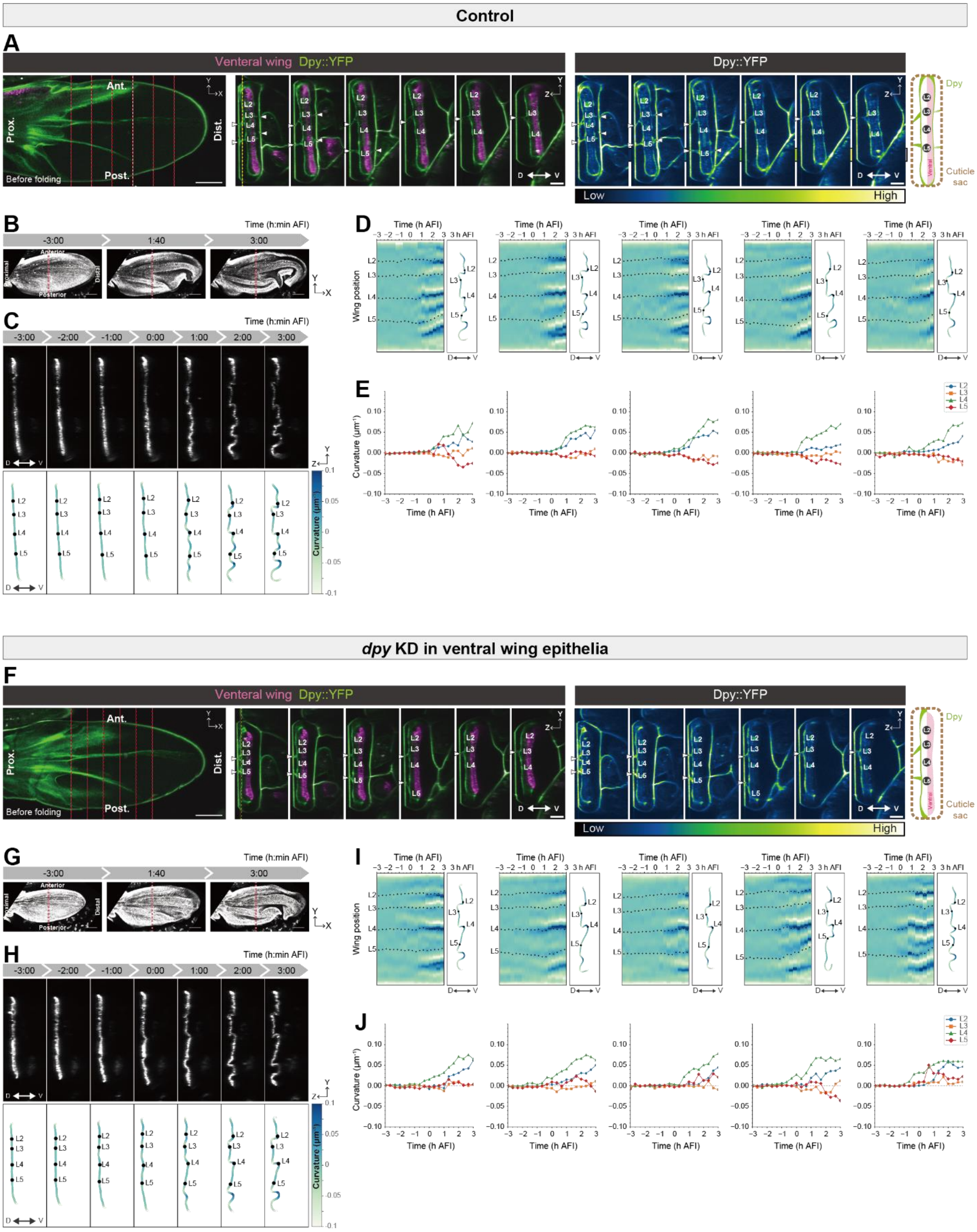


**Fig. S9. Phenotypes of wings with depleted ventral Dpy.**

**(A, F)** MP microscope snapshot images of Dpy::YFP before folding (31:45 APF (A), 32:15 APF (F)) in a control wing (*sd-Gal4>UAS-mCD8::mCherry*, *UAS-Dcr2* and *ap-lexA>LexAop-Gal80*)

(A) and a wing with depleted ventral Dpy (*sd-Gal4>UAS-dpyRNAi, UAS-mCD8::mCherry, UAS-Dcr2,* and *ap-lexA>LexAop-Gal80*) (F). Both wings express mCD8::mCherry to label ventral cells. Left: XY views of a slice between the dorsal cuticle and wing surface. Right: Anterior-posterior cross-sections along the red lines in the left panels. Yellow dotted lines represent the corresponding Z slice position of the left panel. White arrowheads indicate exuvial Dpy along veins between the wing and cuticle. Schematics of representative Dpy (green) distribution on wings labeling ventral cells (magenta) inside a cuticle sac (brown). To genetically manipulate the ventral wing surface, Gal4-UAS system, LexA-lexAop system, and RNA interference (RNAi) were used in combination. We drove the expression of UAS transgenes (UAS-mCD8::mCherry and UAS-Dcr2 for control wings, and UAS-dpyRNAi, UAS-mCD8::mCherry, and UAS-Dcr2 for wings with depleted ventral Dp) throughout the wing pouch under the control of sd-Gal4, while we simultaneously expressed Gal80 (a Gal4 repressor) in dorsal cells by ap-LexA to exclude the expression of UAS transgenes from dorsal cells. This combinatorial approach makes it possible to express UAS transgenes only in the ventral cells, as confirmed by UAS-mCD8::mCherry expression only in ventral cells (magenta in middle panels). In a control experiment that lacks UAS-dpyRNAi transgene, exuvial and surface-covering Dpy is present on both the dorsal and ventral surface (A). In contrast, in an experiment that incorporates UAS-dpyRNAi transgene, exuvial and surface-covering Dpy on the ventral surface was diminished, whereas that on the dorsal surface remained the same as that in the control (F).

**(B–E, G–J)** Confocal time-lapse images and tissue curvature in a control wing (*sd-Gal4>UAS-mCD8::mCherry, UAS-Dcr2* and *ap-lexA>LexAop-Gal80*) (B–E) and wings with depleted ventral Dpy (*sd-Gal4>UAS-dpyRNAi, UAS-mCD8::mCherry, UAS-Dcr2* and *ap-lexA>LexAop-Gal80*) (G–J).

**(B, G)** Maximum projections of confocal time-lapse images. Note that only ventral wing cells are labeled with mCD8::mCherry.

**(C, H)** Anterior-posterior cross-sections along the red dotted lines in (B, G) (top) and their color-coded tissue curvature (bottom). Black dots in the bottom panel represent the positions of L2, L3, L4, and L5 longitudinal veins.

**(D, E, I, J)** Tissue curvature measurement from time-lapse imaging of five wings for each genotype. Each left-most sample is the same as (B, C) and (G, H), respectively. D, I: Spatio-temporal maps of curvature (left). Anterior-posterior cross-section showing tissue curvature at 3 h AFI (right). Black dots represent the positions of L2, L3, L4, and L5 longitudinal veins. E, J: Time evolution of local curvature averaged around longitudinal veins. Note that the wing surface marker in control experiments between Fig. 1D and E and fig. S9D and E is different: In Fig. 1D and E, the dorsal cells are labelled, while in fig. S9D and E, the ventral cells are labelled. For all five ventral Dpy-depleted wings, the buckling direction along the veins was almost the same as that of the control, where the L2 and L4 longitudinal veins localized at the region with positive curvature, and L3 and L5 veins localized at the region with zero to negative curvature.

Scale bars, 100 µm (XY views of A and F, B, G), 50 µm (YZ views of A and F, C, H).

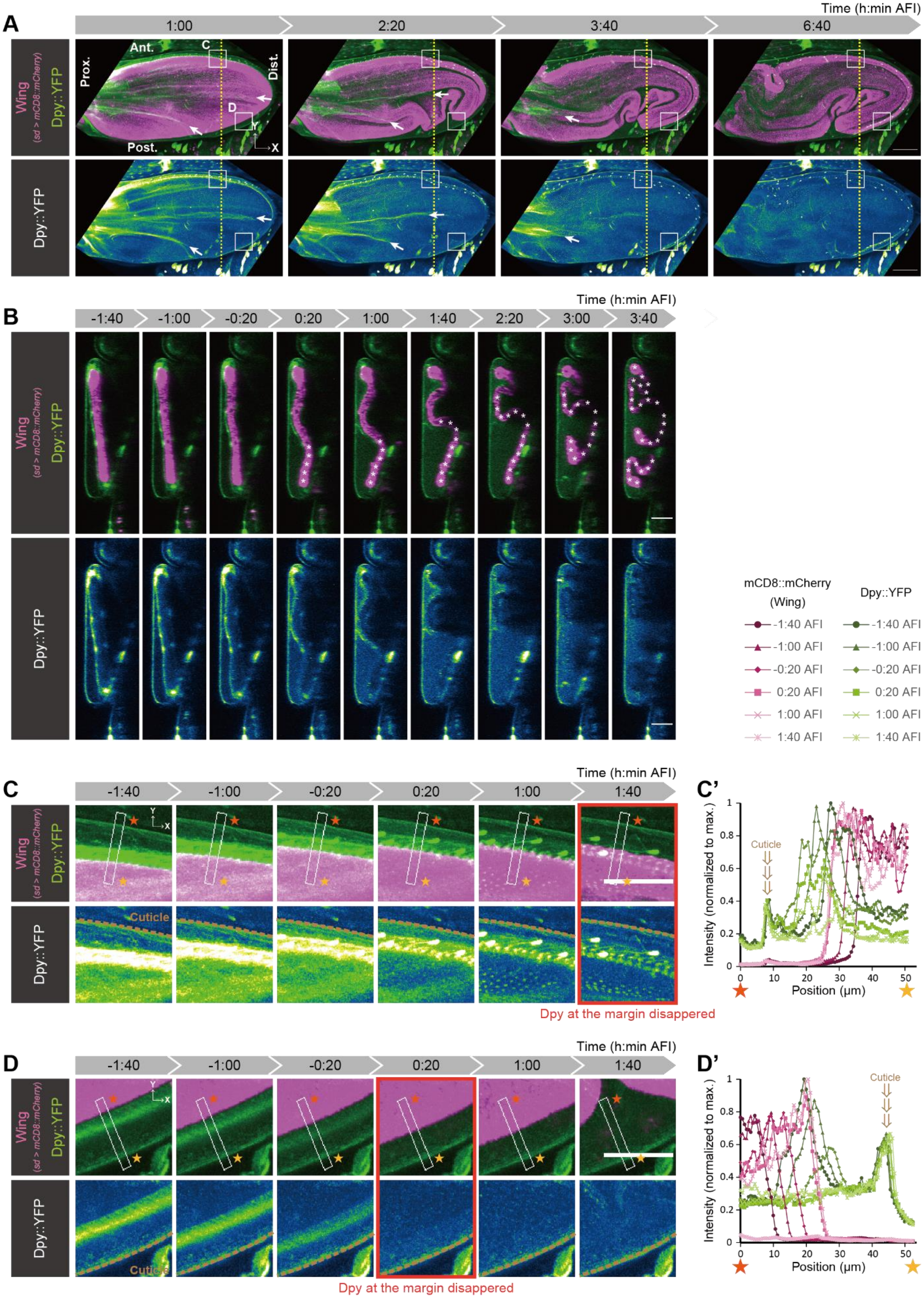
A
Time (h:min AFI)
1:00
2:20
3:40
6:40
Wing (sd > mCD8::mCherry)
Dpy::YFP
Ant.
Post.
Prox.
Dist.
C
D
Y
X
B
-1:40
-1:00
-0:20
0:20
1:00
1:40
2:20
3:00
3:40
mCD8::mCherry (Wing)
Dpy::YFP
-1:40 AFI
-1:00 AFI
-0:20 AFI
0:20 AFI
1:00 AFI
1:40 AFI
C
C'
Cuticle
Intensity (normalized to max.)
Position (μm)
Dpy at the margin disappeared
D
D'

**Fig. S10. Exuvial vein/margin Dpy and surface-covering Dpy disappear during wing folding.**

**(A–D)** Confocal time-lapse images in a control wing expressing mCD8::mCherry induced by sd-Gal4 and endogenous Dpy::YFP. The original image of the wing is the same as Fig. 3E–G, fig. S14A, and fig. S15A and C (top panel).
**(A, C, D)** Maximum projections of the images.
**(A)** Disassembly of exuvial Dpy along L3 and L5 veins (white arrows) proceeds in a distal to proximal direction.
**(B)** Anterior-posterior cross-sections along the yellow dotted line in (A). Asterisks indicate the wing position where Dpy disappeared from the nearby apical surface of the wing tissue.
**(C, D)** Magnified views of rectangles at anterior (C) or posterior (D) positions in (A).
**(C', D')** Intensity profile of mCD8::mCherry and Dpy::YFP along a line extending from the orange star to the yellow star in (C, D). The intensity of exuvial Dpy at the wing margin decreased during the wing folding, indicating that exuvial Dpy at the wing margin was disassembled. The decrease in Dpy::YFP intensity at the exuvial margin starts earlier in the posterior region (D) than in the anterior region (C).
Scale bars, 100 µm (A), 50 µm (B, C, D).

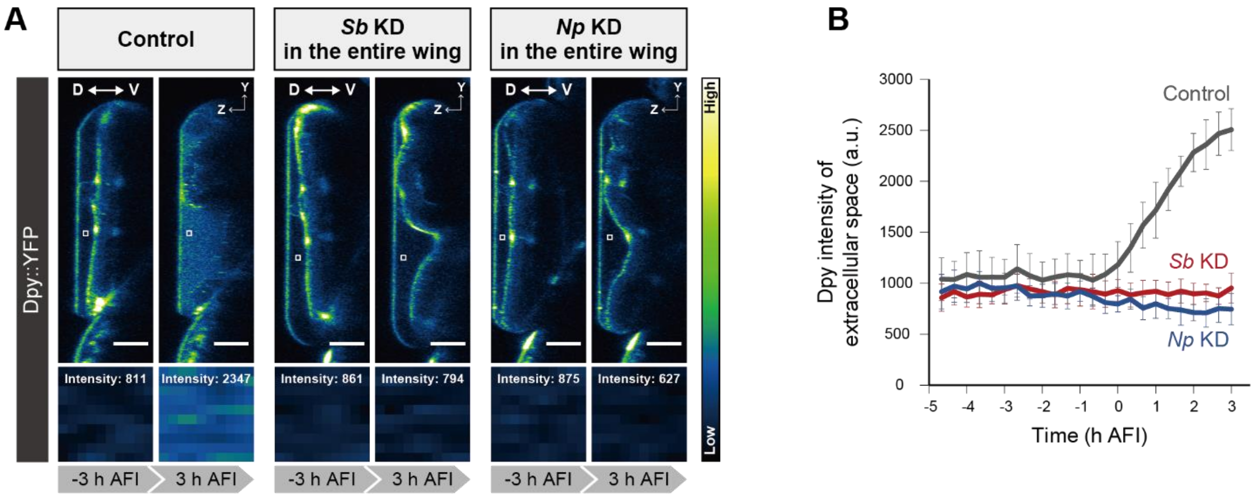


**Fig. S11. Dynamics of Dpy::YFP fluorescent signal at the exuvial space during wing folding.**

**(A)** Anterior-posterior cross-sections of confocal time-lapse images of Dpy::YFP before and after folding (-3 and 3 h AFI) in control (*sd-Gal4>UAS-mCD8::mCherry*), *Sb* knockdown throughout the wing (*sd-Gal4>UAS-mCD8::mCherry, UAS-SbRNAi*), and *Np* knockdown throughout the wing (*sd-Gal4>UAS-mCD8::mCherry, UAS-NpRNAi*). ROIs [10×10 pixels (6.2×6.2 μm)] (white square in top panels) were set at extracellular space between the dorsal cuticle and wing surface to measure the averaged Dpy::YFP intensity in the ROIs (see (B) for the time evolution). The bottom panels show the magnified images of ROIs in the top panels and their averaged intensity.

**(B)** Time evolution (h AFI) of Dpy::YFP intensity in ROIs [10×10 pixels (6.2×6.2 μm)] at extracellular space. N = 5 wings for control, N = 6 wings for *Sb* knockdown, and N = 7 wings for *Np* knockdown. Error bars represent standard deviation.

Scale bars, 50 μm (A).

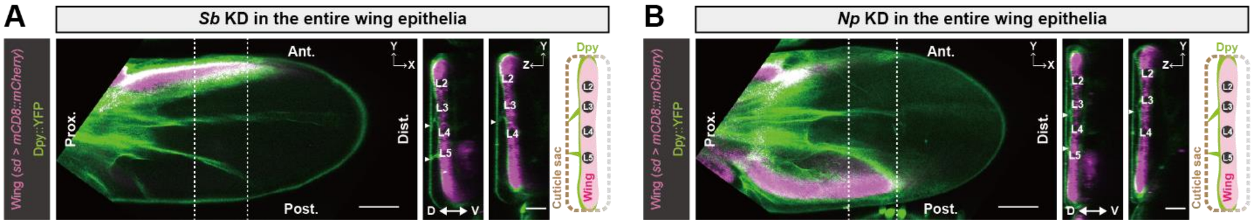


**Fig. S12. Dpy localization in Sb- or Np-depleted wings before folding.**

**(A, B)** Confocal snapshot images of wings expressing mCD8::mCherry induced by sd-Gal4 and endogenous Dpy::YFP before folding (33 h APF) in *Sb* knockdown throughout the wing (*sd-Gal4>UAS-mCD8::mCherry*, *UAS-SbRNAi*) (A) and *Np* knockdown throughout the wing (*sd-Gal4>UAS-mCD8::mCherry*, *UAS-NpRNAi*) (B). Left: XY view of a slice between the dorsal cuticle and wing surface. Right: Anterior-posterior cross-sections along the white dotted lines in the left XY view. White arrowheads indicate the exuvial dorsal Dpy along veins. Schematics showing the representative localization of Dpy in the pupal wing before folding. Green: Dpy, Magenta: Wing, Brown: Surrounding pupal cuticle. L2, L3, L4, and L5 indicate the position of longitudinal veins.

Scale bars, 50 µm (YZ views of A and B), 100 µm (XY views of A and B).

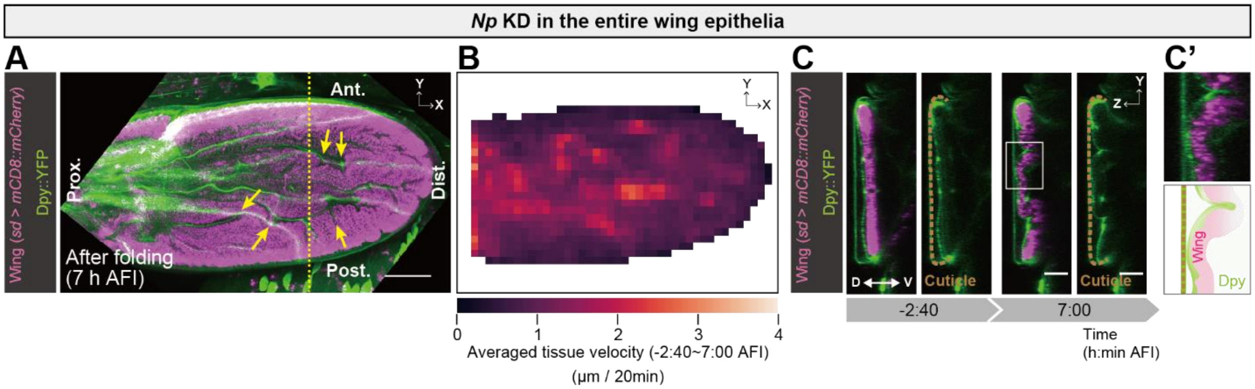


**Fig. S13. Wings with depleted Np failed to undergo Dpy disassembly and to form the marginal fold.**

**(A–C)** Confocal time-lapse imaging of Dpy::YFP and PIV analysis in *Np* knockdown throughout the wing (*sd-Gal4>UAS-mCD8::mCherry, UAS-NpRNAi*). The original image of the wing is the same as fig. S14C and fig. S15A and C (top panel).
**(A)** Maximum projections of post-folded wings (7 h AFI). Yellow arrows indicate folds that cross over the veins.
**(B)** Spatial color maps of tissue velocity averaged in each grid throughout the folding process (-2:40–7:00 AFI).
**(C)** Anterior-posterior cross-sections along the yellow dotted line in (A).
**(C')** Magnified view of the rectangles in (C) and their schematics.
Scale bars, 100 µm (A), 50µm (C).

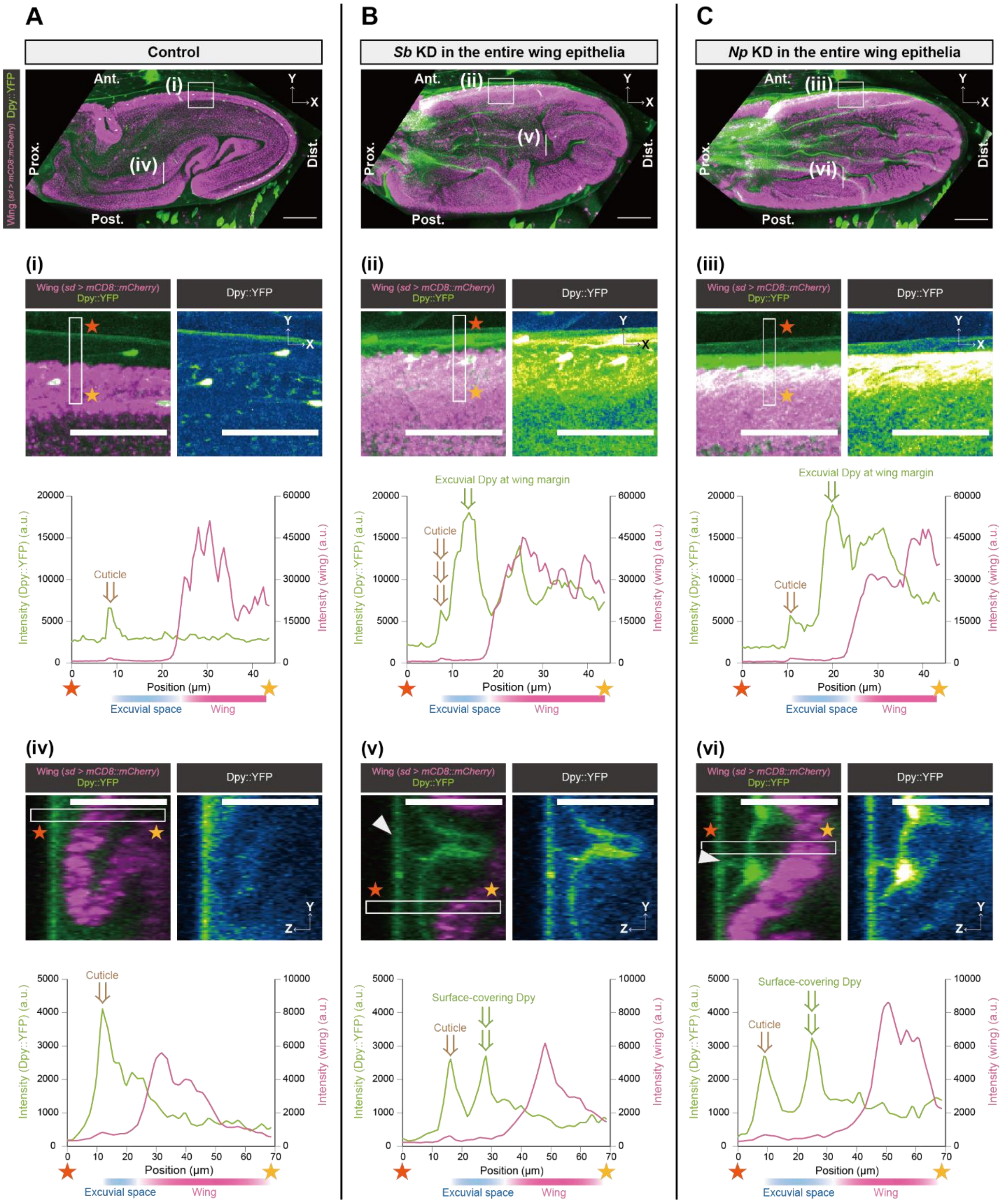


**Fig. S14. Sb and Np depletion prevents disassembly of exuvial and surface-covering Dpy.**
**(A–C)** Top: Maximum projections of post-folded wings (7 h AFI) extracted from confocal time-lapse images of Dpy::YFP in control (*sd-Gal4>UAS-mCD8::mCherry*) (A), *Sb* knockdown throughout the wing (*sd-Gal4>UAS-mCD8::mCherry*, *UAS-SbRNAi*) (B), and *Np* knockdown throughout the wing (*sd-Gal4>UAS-mCD8::mCherry*, *UAS-NpRNAi*) (C). The original image of

the wing is the same as Fig. 3E–G, fig. S10, and fig. S15A and C (top panel) for control, Fig. 3H–J and fig. S15A and C (top panel) for *Sb* knockdown, and fig. S13 and fig. S15A and C (top panel) for *Np* knockdown.

**(i, ii, iii)** Magnified views of rectangles in the top panels and intensity profiles of mCD8::mCherry and Dpy::YFP along a line extending from the orange star to the yellow star. The intensity of exuvial space at the wing margin remained higher in Sb- and Np-depleted wings (see the Dpy::YFP intensity profile at the position indicated by “Exuvial Dpy at wing margin”) compared to control wings, indicating that Sb and Np depletion prevented the disassembly of exuvial Dpy at wing margin.

**(iv, v, vi)** Anterior-posterior cross-sections along the white lines in the top panels. In Sb- and Np-depleted wings, exuvial Dpy along a vein remained in the original position (white arrowheads). Intensity profiles of mCD8::mCherry and Dpy::YFP along a line extending from the orange star to the yellow star show that the intensity of Dpy covering the apical surface of the wings remained higher in Sb- and Np-depleted wings (see the Dpy::YFP intensity profile at the position indicated by “Surface-covering Dpy”) compared to control wings. These results indicate that Sb and Np depletion prevented the disassembly of exuvial vein Dpy and surface-covering Dpy.

Scale bars, 100 µm (top panels), 50 µm (i–vi).

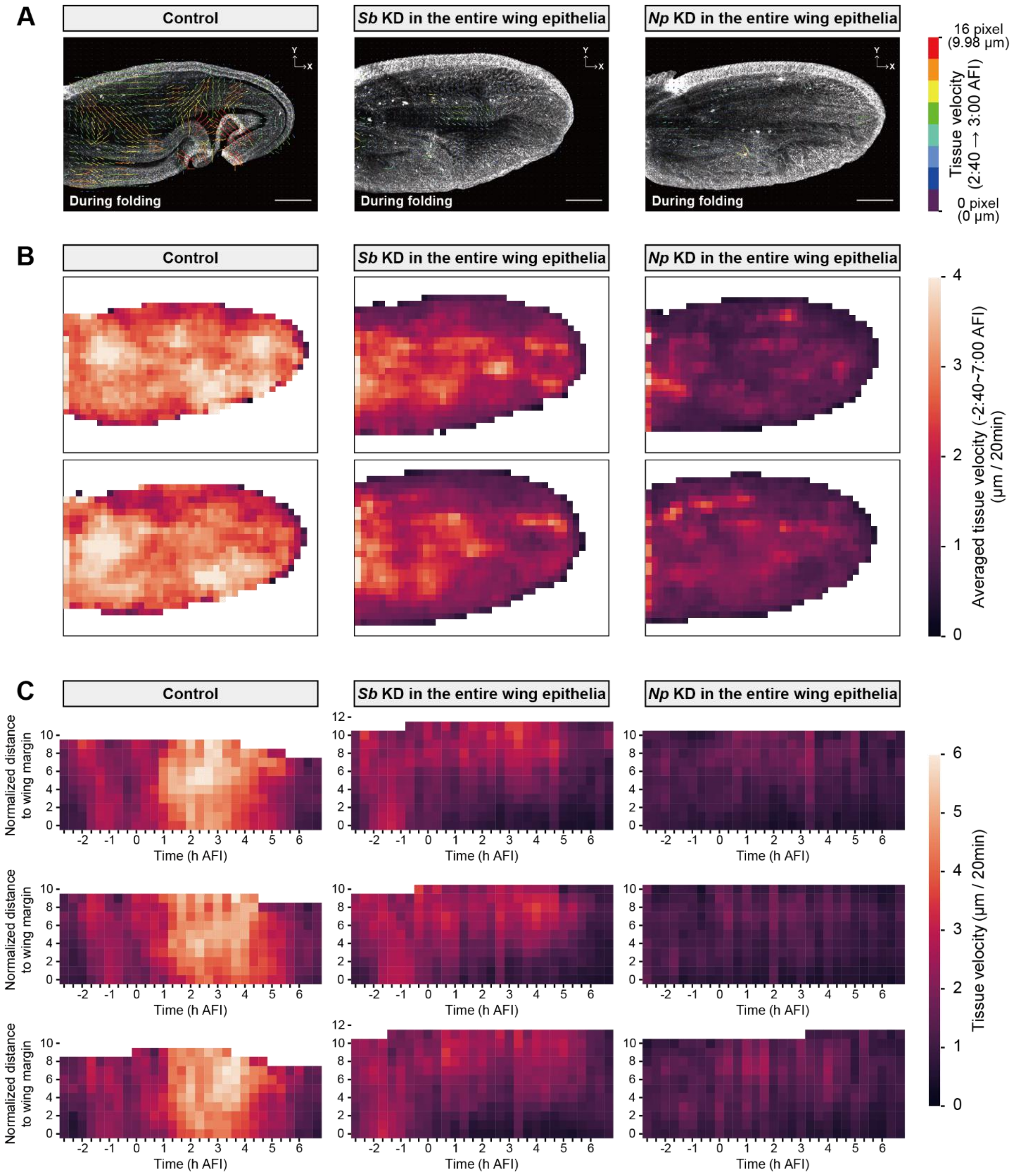


**Fig. S15. Quantification of tissue movement during folding by PIV analysis.**

**(A–C)** PIV analysis in control (*sd-Gal4>UAS-mCD8::mCherry*), *Sb* knockdown throughout the wing (*sd-Gal4>UAS-mCD8::mCherry, UAS-SbRNAi*), and *Np* knockdown throughout the wing (*sd-Gal4>UAS-mCD8::mCherry, UAS-NpRNAi*).

**(A)** Representative examples of tissue movement during folding quantified by PIV analysis. The direction and length of each arrow represent the velocity vectors obtained by the displacement

from 2:40 AFI to 3:00 AFI. Each velocity vector is color-coded according to its magnitude. Original wing images for the PIV analysis are the same as Fig. 3E–G, fig. S10, and fig. S14A for control, Fig. 3H–J and fig. S14B for *Sb* knockdown, and fig. S13 and fig. S14C for *Np* knockdown.
**(B)** Spatial color maps of tissue velocity averaged in each grid throughout the folding process (-2:40–7:00 AFI). Each panel shows individual wing data.
**(C)** Spatio-temporal color maps of tissue velocity magnitude. The X-axis indicates time (h AFI) and Y-axis indicates the normalized minimum distance to the wing margin. Each panel shows the individual wing data (N = 3 wings for each genotype). The top panel of each genotype is generated from the same wing data as Fig. 3E–G, fig. S10, and fig. S14A for control, Fig. 3H–J and fig. S14B for *Sb* knockdown, and fig. S13 and fig. S14C for *Np* knockdown. The middle and bottom panels of each genotype are generated from the same wing data as the top and bottom panels of (B), respectively.
Scale bars, 100 μm (A).

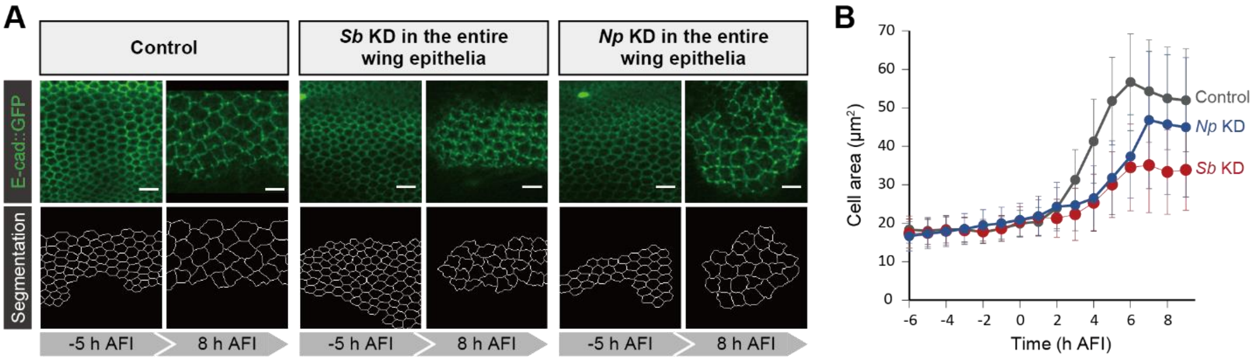


**Fig. S16. Quantification of the apical area of wing epithelial cells in control, *Sb* knockdown, and *Np* knockdown wings during folding.**

**(A, B)** Confocal time-lapse imaging of wings expressing mCD8::mCherry induced by sd-Gal4 and E-Cadherin::GFP before and after wing folding (-5 and 8 h AFI). Control (*sd-Gal4>UAS-mCD8::mCherry*), *Sb* knockdown (*sd-Gal4>UAS-mCD8::mCherry, UAS-SbRNAi*), and *Np* knockdown (*sd-Gal4>UAS-mCD8::mCherry, UAS-NpRNAi*). (A): Maximum projections (top) and segmented images based on junction marker E-Cadherin::GFP (bottom). (B): Cell area as a function of time (h AFI). N = 3 wings (12225 cells) for control, N = 3 wings (11418 cells) for *Sb* knockdown, and N = 3 wings (10411 cells) for *Np* knockdown. Error bars represent standard deviation. In the Sb- or Np-deleted wings, cell area increased gradually, but the area reached after folding (8 h AFI) was smaller than that in control wings, indicating an incomplete cell flattening. Scale bars, 10 μm (A).

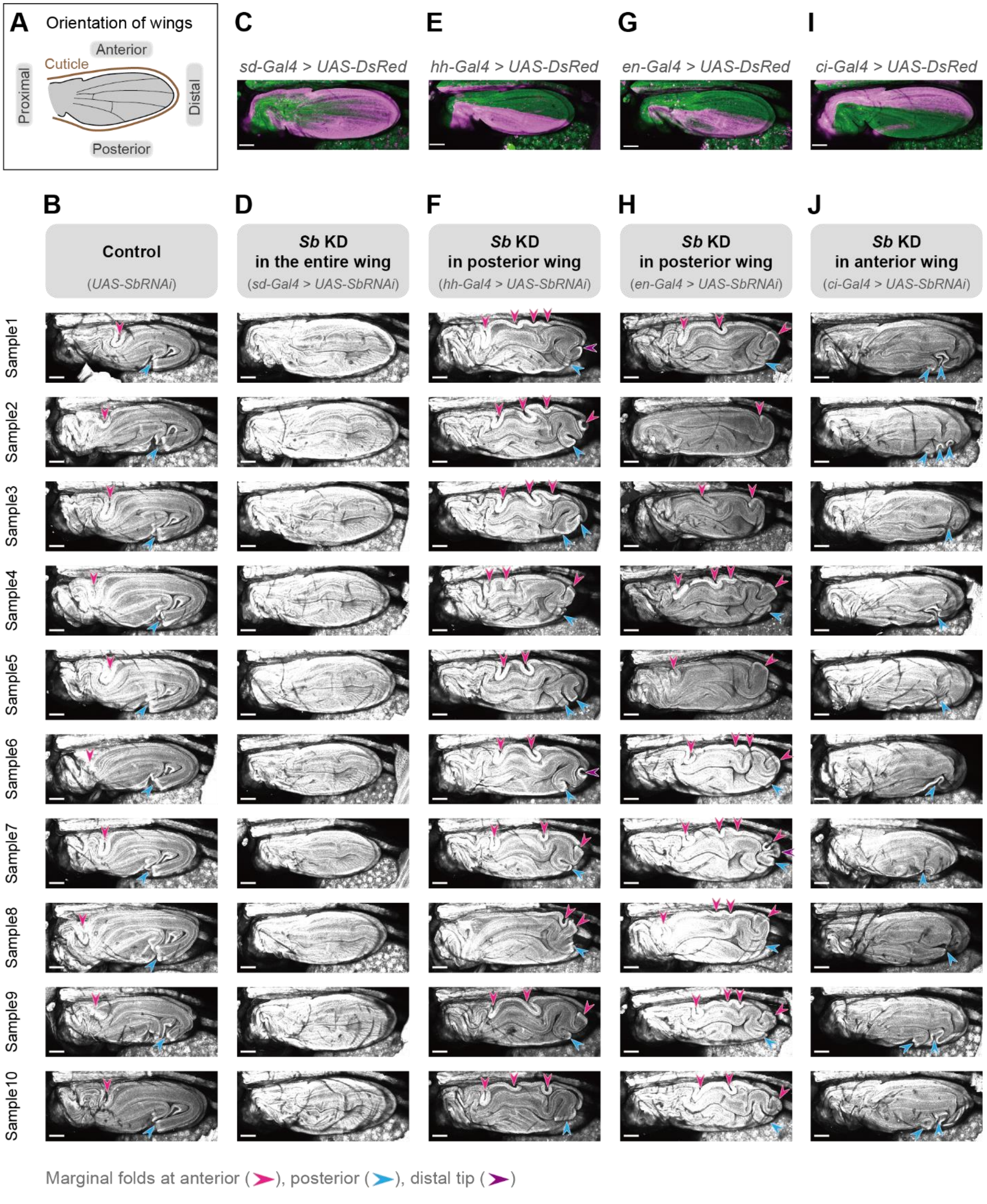


**Fig. S17. Localized silencing of *Sb* with different spatial patterns.**

**(A)** Schematics showing the orientation of wings in this figure.

**(C, E, G, I)** Maximum projections of confocal snapshot images in wings before folding. Wing cells were labeled with GAP43::GFP expressed under the control of ubiquitin promoter (green), and Gal4 expressing regions are marked by UAS-DsRed (magenta). Images of (E, G, I) are the same as fig. S18 C, E, and G.

**(B, D, F, H, J)** Maximum projections of confocal snapshot images of wings after folding in control (*UAS-SbRNAi*) (B) and localized silencing of *Sb* at entire region (*sd-Gal4>UAS-SbRNAi*) (D), posterior region (*hh-Gal4>UAS-SbRNAi* or *en-Gal4>UAS-SbRNAi*) (F, H), and anterior region (*ci-Gal4>UAS-SbRNAi*) (J). Sample dissection for imaging was done after the folding was completed to avoid the artifacts to the marginal fold position associated with dissection before folding (for details, see "Live imaging" section in Materials and Methods). Wing cells were labeled with GAP43::GFP expressed under the control of ubiquitin promoter. Arrowheads represent folds along the wing margin (magenta: anterior, blue: posterior, purple: distal tip). The top three images of B, F, and J are the same as Fig. 4A–A''. Scale bars, 100 μm.

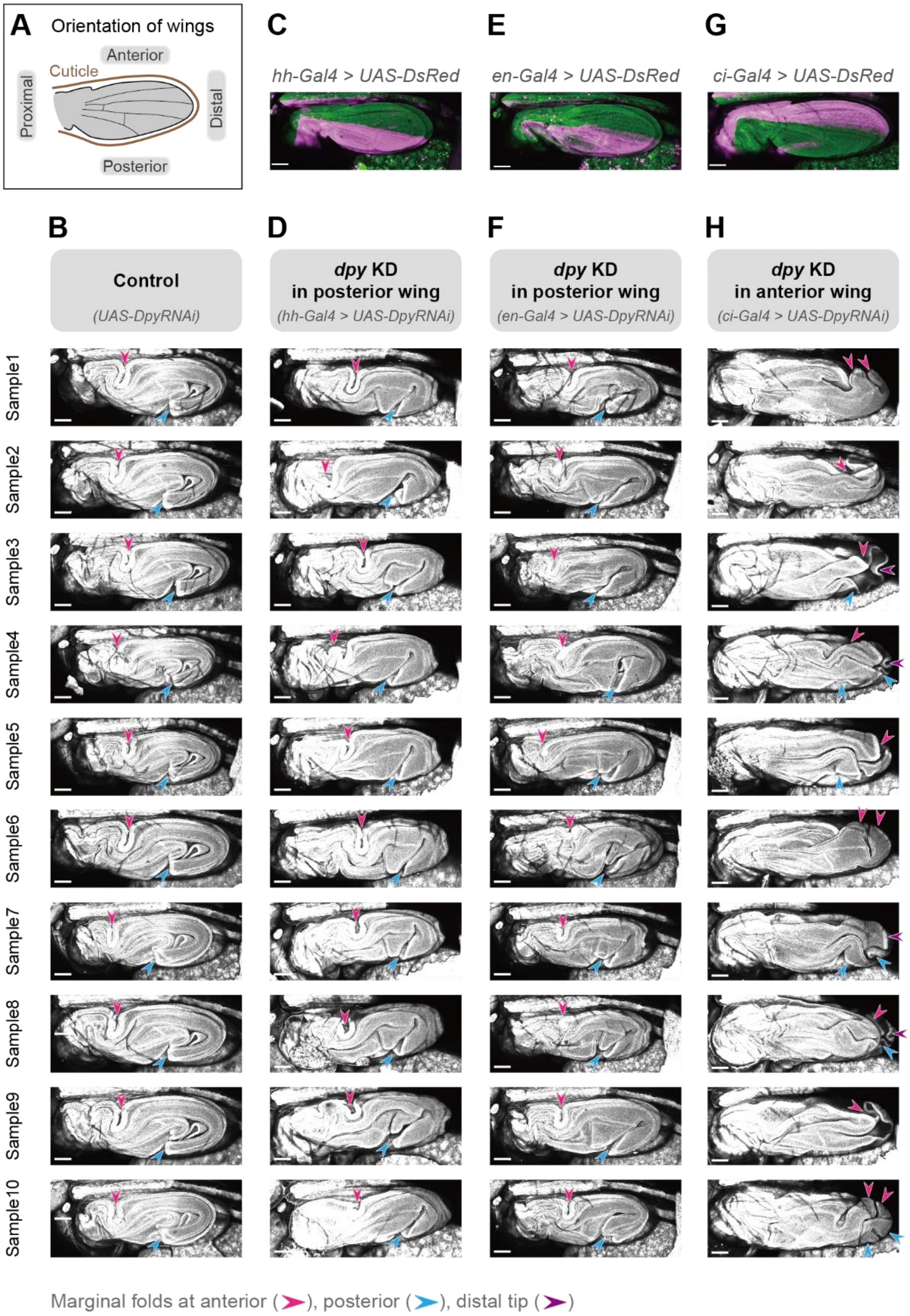


**Fig. S18. Localized silencing of *dpy* with different spatial patterns.**

**(A)** Schematics showing the orientation of wings in this figure.

**(C, E, G)** Maximum projections of confocal snapshot images in wings before folding. Wing cells were labeled with GAP43::GFP expressed under the control of ubiquitin promoter (green), and

Gal4 expressing regions are marked by UAS-DsRed (magenta). Images are the same as in fig. S17E, G, and I.

**(B, D, F, H)** Maximum projections of confocal snapshot images of wings after folding in control (*UAS-dpyRNAi*) (B) and localized silencing of *dpy* at posterior region (*hh-Gal4>UAS-dpyRNAi* or *en-Gal4>UAS-dpyRNAi*) (D, F) and anterior region (*ci-Gal4>UAS-dpyRNAi*) (H). Sample dissection for imaging was done after the folding was completed to avoid the artifacts to the marginal fold position associated with dissection before folding (for details, see "Live imaging" section in Materials and Methods). Wing cells were labeled with GAP43::GFP expressed under the control of ubiquitin promoter. Arrowheads represent folds along the wing margin (magenta: anterior, blue: posterior, purple: distal tip). The top three images of B, D, and H are the same as Fig. 4B–B''.

Scale bars, 100 μm.

**Movie S1.**

Confocal time-lapse imaging of a control wing. Maximum projection (left) and anterior-posterior cross-section along the red solid line in the left panel (right). Time after puparium formation (h:min APF) and after folding initiation (h:min AFI) is shown in the upper-left panel. (Related to Fig. 1A (i) and B).

**Movie S2.**

Maximum projections of confocal time-lapse images in the control wing (top) and their segmented images based on the junction marker E-Cadherin::GFP (bottom). Note that the number of the segmented tracked cells does not change during time-lapse imaging, which indicates that no cell divisions occur during the wing folding process. Time after folding initiation (h:min AFI) is shown in the upper-left panel. Scale bars, 50 µm. (Related to fig. S3B).

**Movie S3.**

Confocal time-lapse imaging of NSlmb-vhhGFP4-mediated knockdown of Sqh::eGFP (green) at anterior region expressing mCD8::mCherry (magenta) induced by ci-Gal4. Maximum projection (left) and anterior-posterior cross-section (right). Time (h:min) after the temperature shift is shown in the upper-left panel. (Related to fig. S5A).

**Movie S4.**

Confocal time-lapse imaging of NSlmb-vhhGFP4-mediated knockdown of Sqh::eGFP (green) at posterior region expressing mCD8::mCherry (magenta) induced by hh-Gal4. Maximum projection (left) and anterior-posterior cross-section (right). Time (h:min) after the temperature shift is shown in the upper-left panel. (Related to fig. S5B).

**Movie S5.**

MP microscope imaging of a control wing expressing mCD8::mCherry (magenta) induced by sd-Gal4 and endogenous Dpy::YFP (green). XY view of a slice at the position of a white line in the right panel (left) and the anterior-posterior cross-sections along white lines in the left panel (right). (Related to Fig. 2A).

**Movie S6.**

Anterior-posterior cross-sections of confocal time-lapse imaging in a control wing expressing mCD8::mCherry (magenta) induced by sd-Gal4 and endogenous Dpy::YFP (green). See the schematic in fig. S7 for the approximate location of cross-sections 1 and 2. Time after folding initiation (h:min AFI) is shown in the upper-left panel. (Related to fig. S7).

**Movie S7.**

Confocal time-lapse imaging of a wing with depleted dorsal Dpy. Maximum projection (left) and anterior-posterior cross-section along the red solid line in the left panel (right). Time after folding initiation (h:min AFI) is shown in the upper-left panel. (Related to Fig. 2D and E).

**Movie S8.**

MP microscope time-lapse imaging of a control wing expressing mCD8::mCherry (magenta) induced by sd-Gal4 and endogenous Dpy::YFP (green). The right and bottom panels show the anterior-posterior and proximal-distal cross-sections along white lines, respectively. The time after folding initiation (h:min AFI) is shown in the upper-left panel. (Related to Fig. 3A–C).

**Movie S9.**

Confocal time-lapse imaging of wings expressing mCD8::mCherry (magenta) induced by sd-Gal4 and endogenous Dpy::YFP (green) in control (upper), *Sb* knockdown (middle), and *Np* knockdown (bottom). Maximum projection (left) and anterior-posterior cross-section (right). Time after folding initiation (h:min AFI) is shown in the upper-left corner. (Related to Fig. 3E, F, F', H, I, and I' and fig. S13A, C, and C').

**Movie S10.**

3D view images rotating around the y-axis in control (upper), *Sb* knockdown (middle), and *Np* knockdown (bottom) wings at 7 h AFI. (Related to Fig. 3E and H and fig. S13A).

**Movie S11.**

Tissue movement during folding quantified by PIV analysis in control (upper), *Sb* knockdown (middle), and *Np* knockdown (bottom). Time after folding initiation (h:min AFI) is shown in the upper-left panel. (Related to fig. S15A).
